# Mavacamten stabilizes the auto-inhibited state of two-headed cardiac myosin

**DOI:** 10.1101/287425

**Authors:** John A. Rohde, David D. Thomas, Joseph M. Muretta

**Affiliations:** Department of Biochemistry, Molecular Biology, and Biophysics University of Minnesota

**Author notes:** Address correspondence to: David D. Thomas, Address: Department of Biochemistry, Molecular Biology, and Biophysics, University of Minnesota, 6-155 Jackson Hall, 321 Church St SE, Minneapolis, MN 55455; Joseph M. Muretta, Address: Department of Biochemistry, Molecular Biology, and Biophysics, University of Minnesota, 6-155 Jackson Hall, 321 Church St SE, Minneapolis, MN 55455.

## Abstract

We used transient biochemical and structural kinetics to elucidate the molecular mechanism of mavacamten, an allosteric cardiac myosin inhibitor and prospective treatment for hypertrophic cardiomyopathy. We find that mavacamten stabilizes an auto-inhibited state of two-headed cardiac myosin, not found in the isolated S1 myosin motor fragment. We determined this by measuring cardiac myosin actin-activated and actin-independent ATPase and single ATP turnover kinetics. A two-headed myosin fragment exhibits distinct auto-inhibited ATP turnover kinetics compared to a single-headed fragment. Mavacamten enhanced this auto-inhibition. It also enhanced auto-inhibition of ADP release. Furthermore, actin changes the structure of the auto-inhibited state by forcing myosin lever-arm rotation. Mavacamten slows this rotation in two-headed myosin but does not prevent it. We conclude that cardiac myosin is regulated in solution by an interaction between its two heads and propose that mavacamten stabilizes this state.

**Significance Statement:** Small-molecule allosteric effectors designed to target and modulate striated and smooth myosin isoforms for the treatment of disease show promise in preclinical and clinical trials. Beta-cardiac myosin is an especially important target, as heart disease remains a primary cause of death in the U.S. One prevalent type of heart disease is hypertrophic cardiomyopathy (HCM), which is hypothesized to result from dysregulated force generation by cardiac myosin. Mavacamten is a potent cardiac myosin ATPase activity inhibitor that improves cardiac output in HCM animal models. Our results show that mavacamten selectively stabilizes a two-head dependent, auto-inhibited state of cardiac myosin in solution. The kinetics and energetics of this state are consistent with the auto-inhibited super-relaxed state, previously only observed in intact sarcomeres.

## Introduction

Familial hypertrophic cardiomyopathies, abbreviated HCM, represent one of the most common classes of genetic disease, affecting 1 in 500 people[1]. HCM is hypothesized to result from cardiac hyper-contractility[2]. Direct inhibition of force generation by cardiac myosin is thus a putative treatment [3]. Mavacamten (Mava), previously termed Myk461, is a small-molecule allosteric inhibitor of cardiac myosin that shows promise in preclinical and clinical trials for the treatment of HCM [3]. Mavacamten binds with sub-micromolar affinity to cardiac myosin in the presence of adenosine triphosphate (ATP) and inhibits steady-state actin-activated and actin-independent (basal) ATPase cycling and calcium regulated ATPase activity in permeablized cardiac myofibrils[3, 4]. Mavacamten also inhibits the kinetics of rigor actin-binding as well as the kinetics of actin-activated phosphate release [4]. It decreases *in vitro* actin filament sliding velocity in the actomyosin motility assay [4], decreases force generation by skinned cardiac preparations from mouse HCM models [3], and decreases cardiac output in live feline hearts [5]. Mavacamten’s effect on cardiac contractility is hypothesized to follow from its inhibition of actin-activated phosphate release, the kinetic step most-tightly coupled to force generation [4].

Because mavacamten changes the kinetics of actin-activated phosphate release, we hypothesized that it may also stabilize a previously un-detected auto-inhibited state of two-headed cardiac myosin in solution, analogous to the super-relaxed state (SRX) observed in skinned myocardium and skeletal muscle fibers[6, 7]. This hypothesis was prompted by studies of blebbistatin, a myosin II inhibitor which also inhibits phosphate release and is proposed to stabilize the SRX state in skeletal and cardiac muscle[8–11].

In permeabilized relaxed muscle and myocardium, the SRX is defined by the bi-exponential kinetics of single ATP turnover. The fast phase of this turnover is similar to ATP turnover in isolated single myosin heads (S1, Fig. 1A), while the slow phase is thought to reflect myosin heads that are auto-inhibited in a structural state that is folded back onto the filament backbone via a direct interaction between the two-heads of the myosin dimer, and the heads and the S2 coiled-coil domain and filament backbone. As a result of these interactions, the folded heads are hypothesized to turn over ATP much more slowly than the unfolded heads or than S1 heads isolated in solution. This is likely because the structural changes in the ATPase site that are required for dissociation of the ATP hydrolysis products require movement of the myosin light-chain binding domain and actin-binding interface[12, 13]. Movement of these elements would be prevented in the folded state due to direct interaction of the heads and the head-S2 domain, and thus, ATP-turnover would be slowed in this state.

**Fig. 1.**
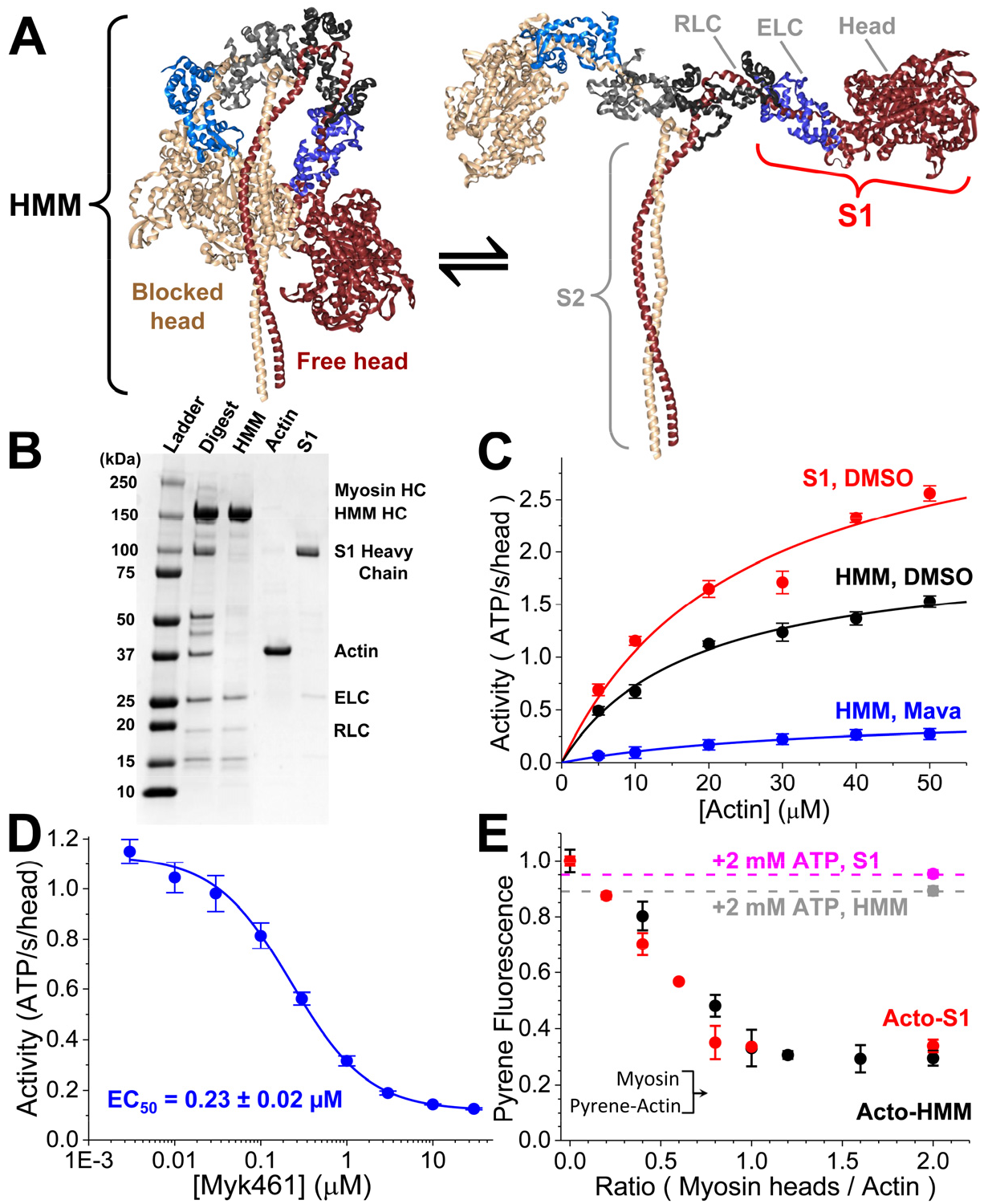
Steady-state ATPase activity of purified cardiac myosin fragments, HMM and S1. (A) Proposed structural isomerization of HMM between a sequestered interacting-heads motif (IHM, PDB: 5TBY, left) and splayed heads (right). (B) SDS-PAGE gel, stained with coomassie, demonstrating the purification of α-chymotryptic-digested HMM and S1 myosin fragments, removing contaminating actin. HC = heavy chain of myosin, ELC = essential light chain, RLC = regulatory light chain. (C) Steady-state, actin-activated ATPase activity of 0.2 µM S1 with DMSO (red), 0.2 μM HMM with DMSO (black) or 10 µM mavacamten (Mava, blue). Activity_S1,DMSO_ = (3.6 ± 0.4 s^−1^)*[Actin]/((24.4 ± 7.4 µM) + [Actin]). Activity_HMM,DMSO_ = (2.02 ± 0.12 s^−1^)*[Actin]/((17.6 ± 2.8 µM) + [Actin]). Activity_HMM,Mava_ = (0.48 ± 0.04 s^−1^)*[Actin]/((35.3 ± 6.3 µM) + [Actin]). Replicates of n=6 for each actin concentration ± SEM. (D) Mavacamten used in this study is a potent inhibitor of 0.2 µM HMM’s actin-activated, steady-state ATPase cycling, [Actin] = 20 µM. Replicates of n=4 ± SEM. Data is not normalized. (E) Varied [S1] from 0-2.0 μM, and varied [HMM] from 0-1.0 µM (0-2.0 μM heads) mixed with 1.0 µM pyrene-labeled actin. Final concentrations listed. The HMM and S1 used in this study releases from pyrene-labeled actin with addition of 2.0 mM ATP, F_S1_ = 0.95 ± 0.1 (magenta), F_HMM_ = 0.89 ± 0.02 (grey). Replicates of n=4 ± SEM. All experiments performed in 10 mM Tris, pH 7.5 at 25 °C, 2 mM MgCl_2_, and 1.0 mM (1,4-dithiothreitol) DTT, unless otherwise noted.

The interactions between myosin heads and the S2 coiled-coil domain and thick-filament backbone described above are collectively termed the interacting-heads motif (IHM, Fig. 1). The IHM is hypothesized to be an evolutionarily conserved mechanism for controlling myosin II driven motility [13–16]. Though electron microscopy (EM) studies suggest that the individual myosin heads in the cardiac thick-filament can form the IHM state [10, 17] it has not been observed in cardiac myosin in solution without chemical stabilization, nor have its associated transient biochemical and structural kinetics been fully explored.

We tested the hypothesis that cardiac myosin is auto-inhibited by direct head-head interaction and that mavacamten selectively targets this intrinsic inhibition by comparing the actin-activated and non-activated (basal) single ATP turnover kinetics of single and two-headed bovine ventricular cardiac myosin fragments. Based on the results from those experiments, we evaluated the temperature and ionic strength dependence of ATP turnover by these same myosin preparations. We reasoned that if the two-headed heavy meromyosin (HMM) forms an IHM-driven, auto-inhibited state, then the energetics of ATP turnover by HMM should reflect this formation and should be much more dependent on increasing ionic strength than S1[18]. Furthermore, if mavacamten stabilizes an auto-inhibited state, it should reduce the state’s sensitivity to increasing ionic strength.

These experiments reveal key aspects of cardiac myosin function and of mavacamten’s mode of action *in vitro*. Foremost, we find that the compound stabilizes a kinetic state of two-headed cardiac HMM that is present to a much smaller degree in single-headed S1. This state is directly observable in the transient kinetics of single ATP turnover by purified cardiac HMM in the absence of mavacamten and reflects the formation of a stable protein-protein interaction interface, indicated by its stability with increasing temperature and by its disruption with increasing ionic strength. Interestingly, actin binding initiates structural changes involving lever-arm rotation of myosin in the auto-inhibited state and this disrupts auto-inhibition. These changes are followed by the actin-activated release of ATP hydrolysis products. Mavacamten slows but does not prevent actin activation of the auto-inhibited state, indicating that it not only stabilizes the state, but that it also increases the transition state free energy required to disrupt it.

## RESULTS

### The steady-state, actin-activated ATPase activity is lower for two-headed cardiac HMM than for single-headed myosin

We prepared soluble myosin fragments for this study as described in our prior work [19] by isolating cardiac myosin from the left ventricles of bovine hearts, digesting the isolated myosin with α-chymotrypsin (Fig. 1B, Lane 2) and purifying the soluble, digested fragments by both size-exclusion and anion-exchange FPLC as described in the SI. Fractions containing two-headed heavy meromyosin (HMM) or the single-headed S1 fragment lacking the regulatory light-chain binding domain and the S2 coiled-coil domain were separated and dialyzed into assay buffers as indicated for each experiment (Fig. 1B, Lanes 3,5). We verified that the purified HMM and S1 were fully intact and did not contain contaminating cardiac actin by gel electrophoresis (Fig. 1B, Lane 4).

We evaluated the functional activity of the purified myosin fragments by measuring actin-activated ATPase activity over a range of actin concentrations using the NADH-coupled ATPase assay[20] (Fig. 1C). Single hyperbolic fits to the actin concentration-dependence of ATPase activity showed that the k_cat_ for the S1 fragment is 3.6 ± 0.4 s^−1^ compared to 2.02 ± 0.12 s^−1^ for cardiac HMM from the identical preparation (p ≤ 0.0035, throughout this manuscript, measurement statistics and statistical significance are tabulated in Table S1). Thus, dimerization of the myosin heads reduces the maximum rate of ATPase cycling, indicating that the two heads of cardiac myosin, connected via the S2 coiled-coil domain, are able to interact and affect the kinetics of actin-activation (Fig. 1A,C).

We verified that the mavacamten preparation (synthesized by EAG Laboratories, verified by mass spectrometry and NMR spectroscopy, described in the SI) inhibits cardiac myosin as demonstrated in prior reports[3] (Fig. 1C,D). Mavacamten potently inhibited the steady-state ATPase cycling of HMM, decreasing the k_cat_ from 2.02 ± 0.12 s^−1^ to 0.48 ± 0.04 s^−1^ (p ≤ 0.001) and shifting the K_m_ from 17.6 ± 2.8 µM to 35.3 ± 6.3 µM (p ≤ 0.05). At 20 µM actin, the EC_50_ for mavacamten’s ATPase inhibition was 0.23 ± 0.02 µM compared to the value for S1 of 0.47 ± 0.006 µM reported in the literature [4] (p ≤ 0.001). The maximum inhibition at saturating mavacamten and 20 µM actin was 0.12 ± 0.006 s^−1^, a 10-fold reduction compared to the activity at 20 µM actin in the absence of the compound.

We determined that both purified myosin fragments freely and equivalently bind to actin in the absence of ATP and dissociate from actin upon the addition of 2.0 mM ATP by equilibrating varying concentrations of HMM or S1 myosin with a fixed concentration of pyrene-labeled actin (Fig. 1E). The fluorescence of pyrene-labeled actin is linearly proportional to the binding of individual myosin heads to actin [20]. The fluorescence decreased linearly with increasing [myosin heads]/[actin] equilibration stoichiometry (Fig. 1E), reaching a maximum at 1:1 stoichiometry with a 70% fluorescence quench, as expected [20]. The quench of pyrene fluorescence by HMM or S1 was identical indicating that both heads of the HMM freely bind actin similarly to free S1 heads. The binding is nearly entirely relieved to 0.89 ± 0.02 or 0.95 ± 0.01 respectively, with the addition of 2.0 mM magnesium ATP (MgATP), and the remaining strongly bound myosin heads are consistent with the expected 5-10% duty-ratio of cardiac myosin estimated in other studies [2].

These experiments are critical for the design and interpretation of later studies in this manuscript and so we briefly note their conclusions. First, the ability of actin to activate cardiac HMM is reduced compared to S1, but this reduction does not reflect the inability of heads to attach to actin as indicated by stoichiometric pyrene-quenching. Thus, the reduced actin-activated ATPase activity (k_cat_) seen in HMM suggests the specific kinetics of actin-activation are constrained by the S2 coiled-coil domain and the tethering of the two heads of the HMM. We further conclude that the mavacamten used in this study potently inhibits cardiac myosin, as expected from prior published work, and that the compound’s EC_50_ for inhibition is lower for HMM than for S1, indicating that the mavacamten binds more effectively to the two-headed HMM than to the single-headed S1 fragment. Importantly, the ATP release experiment shows that the S1 and HMM are functional as ATP incubation increases pyrene-actin fluorescence thus demonstrating both S1 and HMM freely dissociate from actin during ATP cycling and that the reduced actin-activation seen in the HMM does not simply reflect “dead” or non-functional myosin heads which do not detach from actin.

### Mavacamten inhibits transient, actin-activated single ATP turnover more for two-headed cardiac HMM than for single-headed S1

We investigated the transient kinetics of actin-activated ATP turnover by cardiac myosin HMM and S1 using fluorescently-labeled ATP, mant-ATP (2’- or 3’-O-[N-methylanthraniloyl] adenosine 5’-triphosphate). We mixed the purified myosin fragments by stopped-flow with fluorescent mant-ATP, aged the sample for 2.0 seconds to allow for ATP binding and hydrolysis, and then mixed the resulting steady-state sample with increasing concentrations of actin in the presence of 2.0 mM MgATP. The fluorescence of mant-ATP is enhanced when bound by myosin, and the dissociation of mant-ADP after a single kinetic cycle results in a fluorescent decay (Fig. 2A,D). We performed experiments with replicates indicated in Fig. 2 and Table S1.

**Fig. 2.**
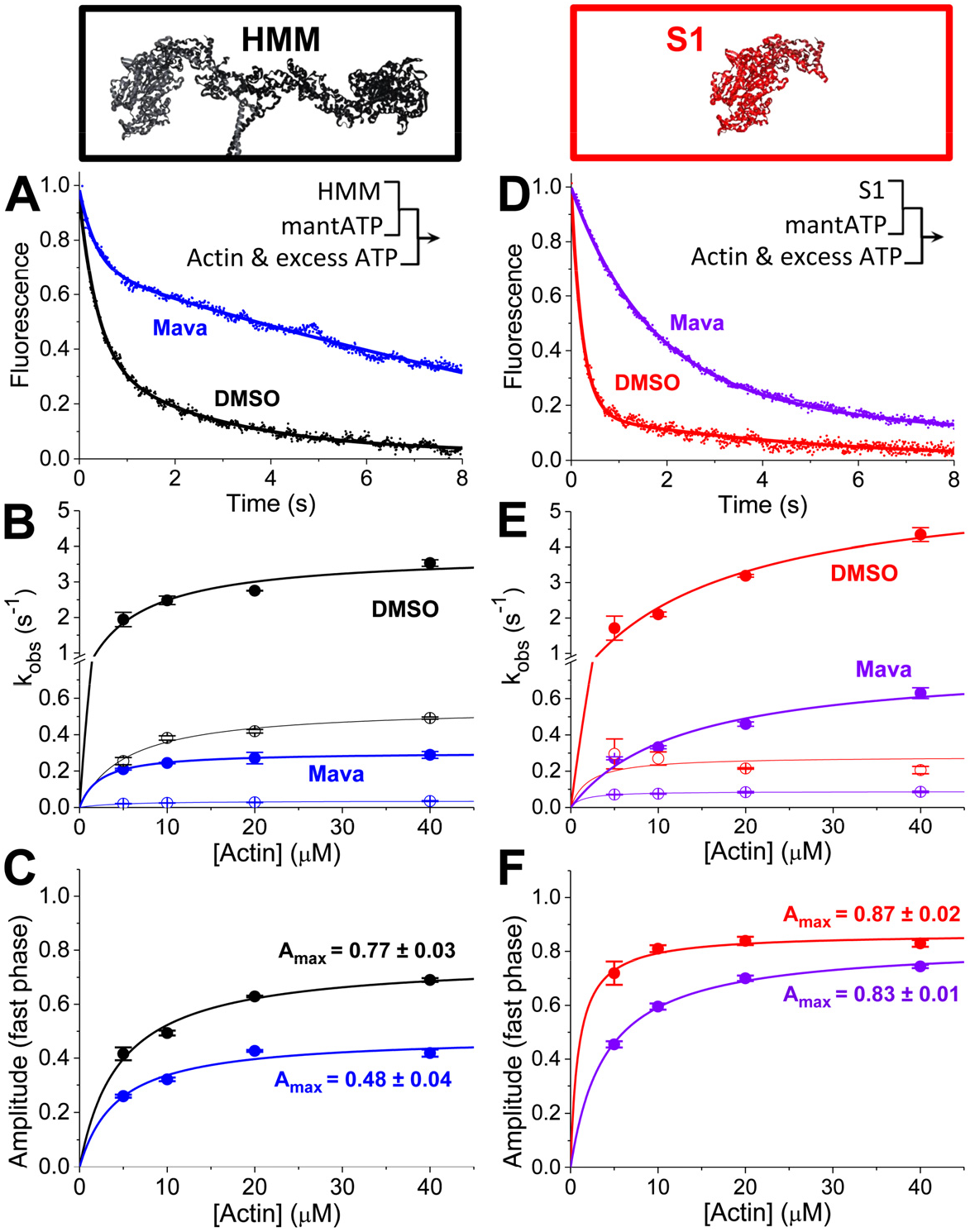
Actin-activated single ATP turnover by (A) 0.1 µM HMM (DMSO control in black, 30 µM mavacamten in blue), mixed with 2.0 μM mant-ATP and 10 μM actin and 2.0 mM MgATP by sequential stopped flow. Pre-mix concentrations listed. Sequential stopped-flow mix schematic is inset. (B-C) Two-exponential fits showing rates and amplitudes with varied [actin]. HMM data is shown in the left column of plots; S1 data is shown in the right column of plots. (D) 0.2 µM S1 (DMSO in red, 30 µM mavacamten in violet), mixed with 2.0 μM mant-ATP and 10 μM actin and 2.0 mM MgATP by sequential stopped flow. (E-F) Two exponential fits showing rates and amplitudes, with varied [actin]. Replicates of n=6 ± SEM, Table S1.

The fluorescent transients were best fit by a bi-exponential time-dependent function consistent with previous published studies of actin-activated ATP turnover by cardiac myosin preparations, Fluorescence = Afast⋅exp(−k_fast_⋅t) + A_slow_⋅exp(−k_slow_⋅t) [21]. We evaluated the actin dependence of the two rate constants (k_fast_ and k_slow_) and the normalized amplitudes (A_fast_ and A_slow_) for each phase in the presence or absence of mavacamten (Fig. 2). The fast phase (closed circles) reflects the release of ATP from a primed, post-hydrolysis, myosin.ADP.P_i_ state ready to undergo actin-activation, while the slow phase (open circles) reflects actin-dependent release from states which are activated more slowly [19, 21]. Importantly, the fast and slow phases are both faster than basal single ATP turnover in the absence of actin (Fig. 3), and as indicated by their increase with increasing actin concentration. The maximum observed rate constant for the fast phase of transient actin-activated ATP turnover was larger for S1 (5.97 ± 0.80 s^−1^) than for HMM (3.80 ± 0.33 s^−1^, p < 0.05), consistent with the steady-state actin-activated ATPase measurements in Fig. 1. Mavacamten inhibited the maximum observed rate constant for the fast phase of actin-activated ATP turnover to 0.80 ± 0.11 s^−1^ in S1 and 0.30 ± 0.01 s^−1^ in HMM. This 7-to 12-fold reduction is similar to the 10-fold inhibition of actin-activated steady-state ATPase cycling (Fig. 1D). The maximum observed rate constant for the slow phase was reduced from 0.28 ± 0.04 s^−1^ to 0.089 ± 0.002 s^−1^ in S1 (a 3-fold inhibition), and from 0.55 ± 0.03 s^−1^ to 0.04 ± 0.01 in HMM (a 14-fold inhibition). Thus, in HMM the slow phase of ATP turnover is significantly more inhibited than in S1 (p ≤ 0.0001). In the absence of mavacamten, the amplitude of the slow phase of actin-activated single ATP turnover is larger in HMM than in S1 (Fig. 2C,F). At saturating actin concentrations, mavacamten decreased the amplitude of the fast phase in HMM (0.48 ± 0.04) but not in S1 (0.83 ± 0.01, p ≤ 0.0001, S1 compared to HMM, Fig. 2C,F). The primary difference between S1 and HMM is the presence of the RLCs and S2 coiled-coil domain, and the dimerization of two myosin heads (Fig. 1A). This dimerization allows the two heads to directly and allosterically interact with each other. Head-head interactions play critical roles in regulating actin-activation in other myosin II family members[13]. The selective effect of mavacamten on the amplitudes in HMM but not S1 at saturating concentrations of actin indicates that mavacamten stabilizes a unique state in HMM, which occurs infrequently in S1 when the myosin heavy chain fragment lacks a regulatory light chain (RLC) and is not tethered to a second myosin by the S2 coiled-coil domain.

**Fig. 3.**
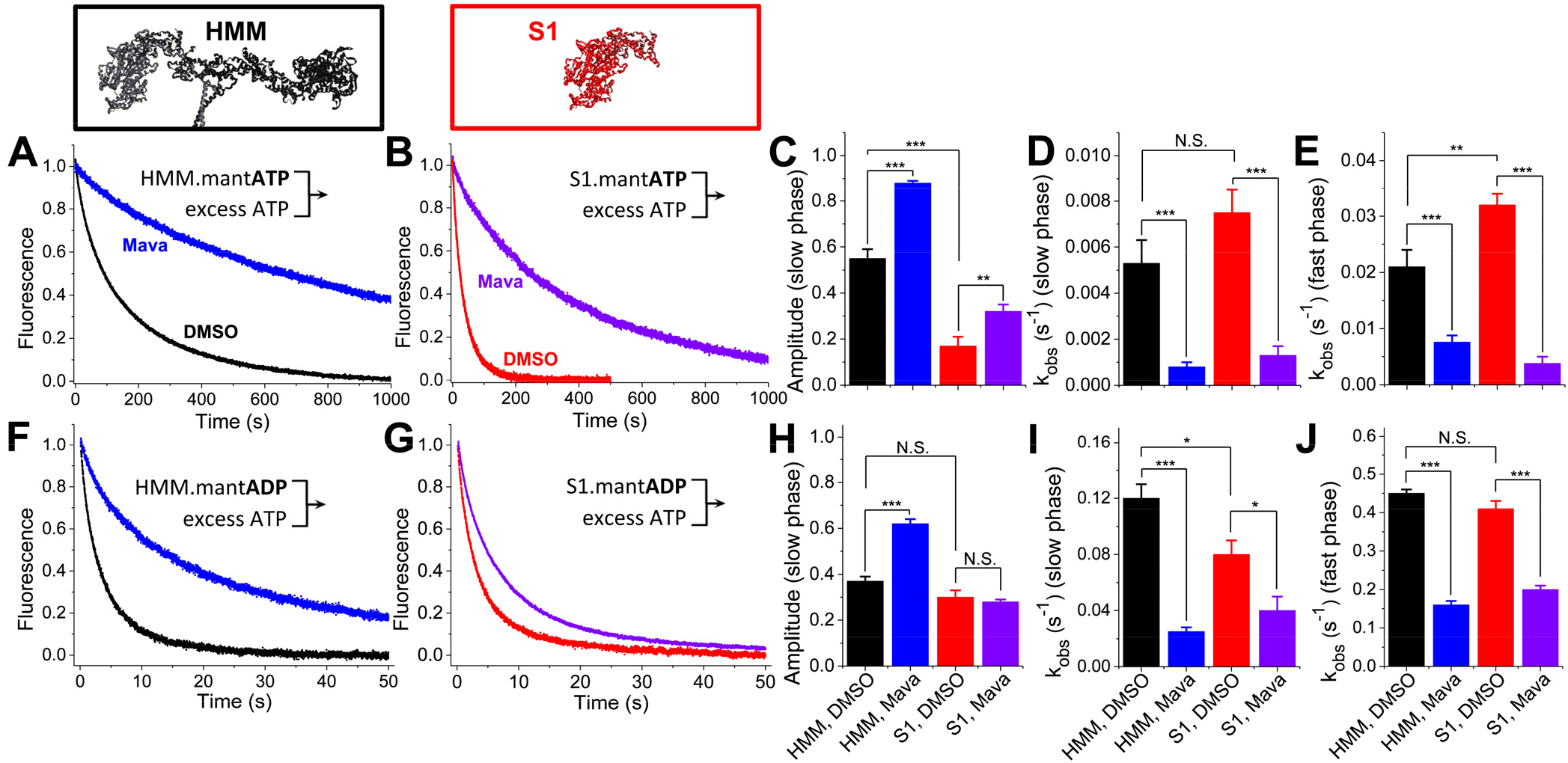
Basal nucleotide exchange in the absence of actin. (A) Basal single ATP turnover by stopped-flow mix of 0.2 µM HMM or (B) 0.4 µM S1, mixed with 4.0 μM mant-ATP, then chased with 2.0 mM MgATP. Blue and violet traces indicate 30 µM mavacamten. (C-E) Amplitudes and rates of the two-exponential fits of A and B. (F) Basal single ADP dissociation from HMM or (G) S1, mixed with 4.0 μM mant-ADP then chased with 2.0 mM MgATP. (H-J) Amplitudes and rates of the two-exponential fit of F and G. Replicates of n = 9, ± SEM. Two-exponential fits reported in Table S1.

### Mavacamten inhibits basal single ATP turnover and basal ADP release in two-headed cardiac HMM differently than in single-headed S1

We investigated the differences between S1 and HMM in the absence of actin by examining the effects of mavacamten on the kinetics of basal single ATP turnover and basal ADP release. We mixed 0.4 μM S1 or 0.2 μM HMM—identical concentrations of ATP-binding heads—with either 4.0 μM mant-ATP (Fig. 3A-E) or 4.0 μM mant-ADP (Fig. 3F-J) and then measured nucleotide release after mixing with 2.0 mM MgATP. Mavacamten’s effect on ADP release was previously measured in S1 but not in HMM (Fig. 3G)[4].

We analyzed the kinetics of nucleotide exchange in these experiments by fitting the data to a bi-exponential function. Basal ATP turnover by HMM was distinct from the ATP turnover kinetics measured in S1 (Fig. 3A-C, black and red traces and bars) exhibiting two distinct kinetic phases. HMM’s slow phase has an amplitude of 0.55 ± 0.04, significantly more than in S1, 0.17 ± 0.04 (Fig. 3C). The rate constant for the fast and slow phase of basal single ATP turnover are very similar in HMM and S1 (Fig. 3 D, E, I, J: compare black and red bars). The rate constants we observed for the fast and slow phases of basal single ATP turnover are 0.02-0.03 s^−1^ and 0.005-0.008 s^−1^ respectively (Table S1), notably similar to the rate constants for bi-exponential mant-ATP turnover kinetics detected in permeablized myocardium where the auto-inhibited SRX state has been described [7].

Mavacamten inhibited the rate constants for basal single ATP turnover in both HMM and S1 preparations to a similar degree (Fig. 3D, E, I, J: blue and violet bars). However, as with the actin-activated single ATP turnover experiments in Fig. 2, the amplitude of the slow phase of basal single ATP turnover is significantly increased by mavacamten in HMM, more than in S1 (Fig. 3C: compare black and blue to red and violet) (p ≤ 0.0001).

The kinetics of ADP release are very similar in HMM and S1 in the absence of mavacamten (Fig. 3F-J: compare black and red traces and bars). As with ATP turnover, ADP release is inhibited more in HMM (Fig. 3F,G: compare blue and violet traces). This increased inhibition reflects stabilization of a slow phase for ATP dissociation that is greatly enhanced from a mole fraction of 0.37 ± 0.02 to 0.62 ± 0.02 (Fig. 3F,H: compare black and blue traces and bars). The fast and slow rate constants for ADP release are similar in S1 and HMM (Fig. 3I,J: black and red bars). But unlike in HMM, mavacamten does not affect the amplitude of the slow phase in S1 (Fig. 3H, red, violet). These results further support the hypothesis that the structural kinetics of HMM are distinct from S1, and mavacamten stabilizes the slow phase of nucleotide turnover. The approximately 20% amplitude of a slow phase in S1 in the absence of mavacamten may reflect a state that is stabilized by protein-protein interactions that occur in two-headed myosin when the heads are tethered, but thermodynamically is still present in a single head to a lesser extent.

### The energetics of basal single ATP turnover in two-headed cardiac HMM are distinct from that of single-headed S1

We evaluated the biophysical determinants underlying the difference between S1 and HMM described in Fig. 3 by examining the temperature dependence of basal single ATP turnover in the absence of actin and in the absence of mavacamten. We performed these experiments identically to those in Fig. 3 over a range of temperatures (5.0 to 35 °C), analyzing the resulting basal single ATP turnover transients by fitting a bi-exponential function to the data. The results from these experiments are summarized in Fig. 4. The kinetics of basal single ATP turnover were distinctly bi-exponential above 15 °C; we focused our analysis above this temperature (Fig. 3B,C,E,F). The rate constants for the fast phase of ATP turnover increased with increasing temperature in both HMM and S1 and the temperature dependence for the increase in k_fast_ and k_slow_ was nearly identical for the two proteins (Table S1). Fitting to the Erying equation provides transition state enthalpy (ΔH^ǂ^) and a collection of terms associated with the transition state entropy (ΔS^ǂ^), equation given in Table S1. For both HMM and S1, the fast rate constant of basal single ATP turnover (k_fast_) has a larger, endothermic ΔHǂ than the slow rate constant (Table S1). In addition, for both HMM and S1, k_fast_ has a positive, dissociative ΔSǂ, while the k_slow_ has a negative, associative ΔSǂ. Fitting the temperature dependence to the Erying equation suggests that k_fast_ and k_slow_ (Fig. 4B, E) are distinct biophysical processes exhibiting unique transition state energetics and are rate-limited by unique biochemical or biophysical transitions. The temperature dependence for the amplitudes of basal single ATP turnover (Fig. 4C,F) were also distinct between S1 and HMM samples. In S1, increasing temperature from 20 to 35 °C dramatically decreased the amplitude of k_slow_ from 0.45 ± 0.07 to 0.07 ± 0.03 (Fig. 4F) while in HMM the amplitude only decreased from 0.57 ± 0.01 to 0.51 ± 0.02 (Fig. 4C). Thus, near physiologic temperatures (35 °C) the slow phase of basal single ATP turnover is significantly more abundant in HMM than in S1 (p ≤ 0.0001). Temperature stability is consistent with the biochemically sequestered SRX state observed in permeablized muscle fibers [6], and the ordered state of the myosin thick-filament observed by fluorescence polarization [22], and X-ray diffraction [23–25].

**Fig. 4.**
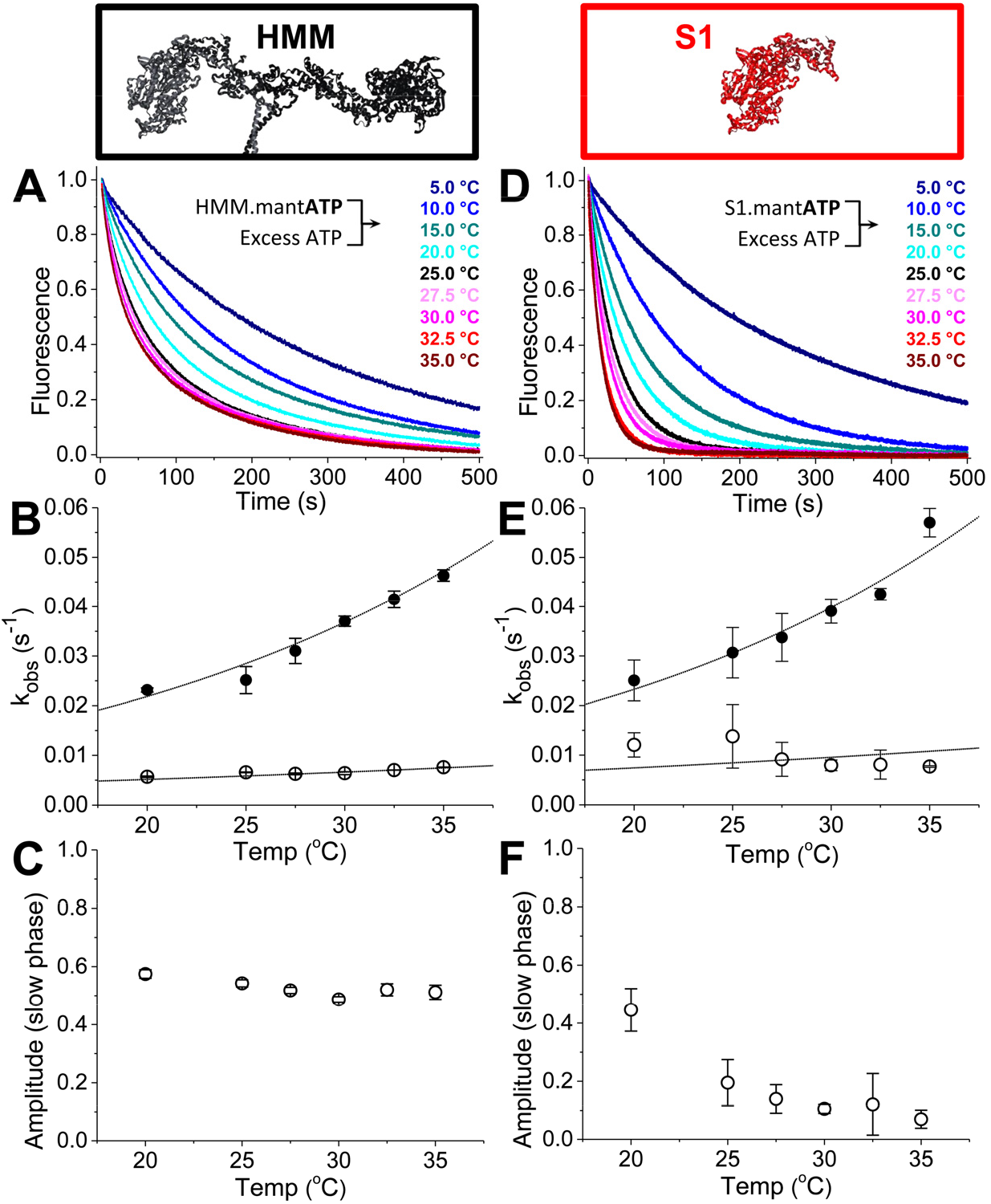
Temperature dependence of basal single ATP turnover. (A) 0.4 µM HMM, or (D) 0.8 µM S1, mixed with 8.0 µM mant-ATP, then mixed with 2.0 mM MgATP. These data were fit to a two-exponential function. (B) The rates for HMM were fit to the Erying equation: k_fast_ = T*exp(–[35.3 kJ/mol]/RT + 5.3) s^−1^K^−1^, k_slow_ = T*exp(–[15.9 kJ/mol]/RT – 4.3) s^−1^K^−1^, with temperature T in Kelvin. Fast phase closed circles; slow phase open circles. (C) The amplitudes of the slow phases depicted in A. (E) The rates from the two-exponential fit for S1 were also fit to the Erying equation: k_fast_ = T*exp(– [30.9 kJ/mol]/RT + 3.6) s^−1^K^−1^, k_slow_ = T*exp(–[15.1 kJ/mol]/RT – 3.9) s^−1^K^−1^. (F) The amplitudes of the slow phases depicted in D. Replicates of n = 4 for each temperature, ± SEM.

### Increasing ionic strength activates two-headed cardiac HMM but not S1

Previous studies showed an isolated S2 coiled-coil domain fragment of human cardiac myosin interacts at low-ionic strength with an isolated S1 fragment and with a two-headed HMM truncated at the second heptad of the S2 coiled-coil domain; and an EM study showed the IHM is disrupted in myosin II homologues by increasing ionic strength [13, 18]. We therefore examined the ionic strength dependence of basal single ATP turnover in S1 and HMM at 25 °C increasing the concentration of potassium chloride (KCl) from 0 mM to 100 mM. The results from these experiments are depicted in Fig. 5. Increasing ionic strength accelerated basal ATP turnover by HMM (Fig. 5A-C) but had little effect on S1 (Fig. 5E-G). We fit bi-exponential functions to the resulting transients. The k_fast_ and k_slow_ rate constants for basal ATP turnover by HMM did not change with increasing ionic strength (Fig. 5B), similar to S1 (Fig. 5F). Furthermore, in S1, the amplitudes of the fast and slow phases also did not change with increasing KCl (Fig. 5G). Conversely, in HMM, increasing ionic strength caused a significant increase in the amplitude of the fast phase and a corresponding decrease in the amplitude of the slow phase of basal single ATP turnover. At 100 mM KCl, these amplitudes were similar to S1 (Fig. 5C,G). These data suggest that the transition between fast- and slow-ATP turnover kinetics states exchange in solution, and that the energetics of this exchange are dependent on charge-charge interactions that are disrupted with increasing ionic strength.

**Fig. 5.**
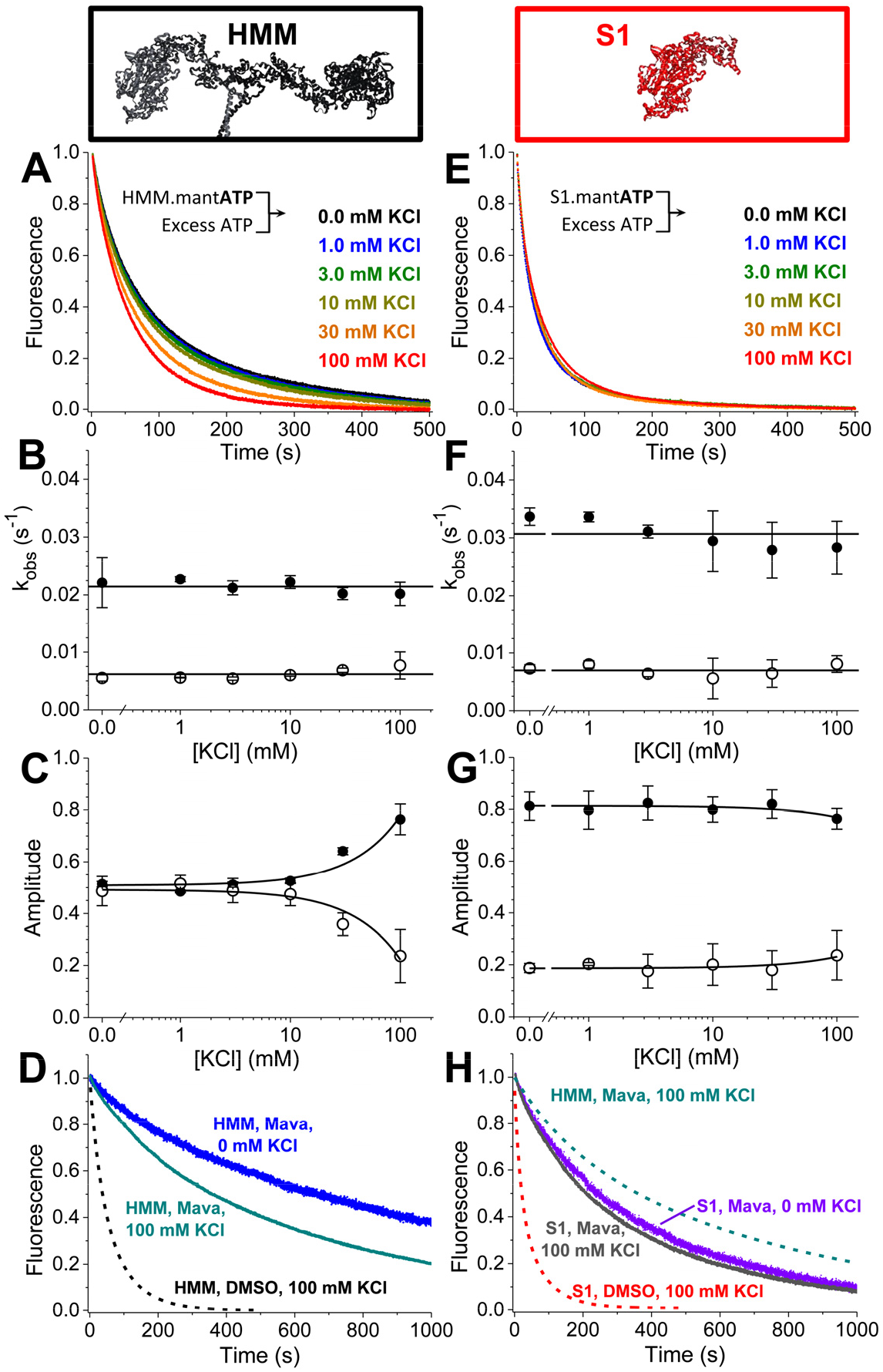
Ionic strength dependence of basal ATP turnover. (A) 0.4 µM HMM, or (E) 0.8 µM S1, mixed with 8.0 µM mant-ATP, then mixed with 2.0 mM MgATP. Data best fit to two exponentials. (B) The rates of HMM’s basal mant-ATP turnover are relatively constant over these [KCl] with average value depicted as a horizontal line. Closed circles represent the fast phase; open circles the slow phase. (C) The amplitude of the fast phase increases with increasing [KCl] (closed circles fast phase; open circles slow phase). Linear fits to show trends. (D) In the presence of mavacamten, HMM is sensitive to increasing ionic strength (blue to dark cyan). (F-G) The rates and amplitudes of S1’s basal mant-ATP turnover are relatively constant. Linear fits to show trends. (H) S1 in the presence of mavacamten is insensitive to changes in [KCl] (violet to grey), consistent with S1 in the absence of mavacamten (E). Replicates of n = 4 for each [KCl], ± SEM. Fits reported in Table S1.

We also examined the ionic strength dependence of HMM’s auto-inhibited state in the presence of mavacamten by performing basal single ATP turnover under 100 mM KCl or no KCl conditions (Fig. 5D). Increasing ionic strength to 100 mM KCl partially relieved mavacamten’s inhibition on ATP turnover in HMM but not in S1, indicating that the mavacamten-stabilized state in HMM is specifically ionic strength dependent (Fig. 5H). Notably, however, at 100 mM KCl and saturating mavacamten, ATP-turnover by HMM is still slower and distinctly bi-exponential than the turnover by S1 (Fig. 5H: compare dashed cyan to violet).

### Actin disrupts the auto-inhibited state of two-headed cardiac HMM

Results in Fig. 3, Fig. 4, and Fig. 5 show that ∼50% of the myosin ATPase sites in HMM turnover ATP much slower than S1, and that this population is stable with increasing temperature and disrupted at physiological ionic strengths. Furthermore, we observed in Fig. 1 and Fig. 2 that the steady-state and transient kinetics of actin-activated ATP turnover in the presence of mavacamten are faster than the kinetics in the absence of actin, depicted in Fig. 3. Taken together, these results show that actin interaction relieves the auto-inhibition present in HMM, even when mavacamten is bound. Furthermore, if mavacamten stabilizes a folded, IHM-like structural state in cardiac myosin, then the structural dynamics of the two light-chain binding domains of the HMM dimer should be altered by the compound. We therefore used transient time-resolved FRET, (TR)^2^FRET, to measure these dynamics directly in response to actin activation.

We performed these experiments using a similar strategy as described in our prior work[19]. We labeled the cardiac myosin regulatory light-chain (RLC) with the AlexaFluor-488 fluorescent probe on a single engineered cysteine residue, replacing the native valine at position 105 in the bovine cardiac RLC sequence. We then exchanged the labeled RLC onto purified cardiac HMM to obtain donor-labeled HMM. Expression and purification of the RLC, labeling, and light-chain exchange were performed identically to our prior study and are described in detail in that paper and outlined in the SI [19]. We confirmed that neither the exchange nor the attachment of the fluorescent labels perturb the auto-inhibited state by measuring basal single ATP turnover (Fig. S2).

We measured actin-initiated changes in light-chain domain (lever arm) orientation by equilibrating the donor labeled HMM with 20-molar excess Cy3-ATP. The Cy3-ATP binds and is hydrolyzed by the myosin. When bound, the Cy3 fluorescent probe is a FRET acceptor to the Alexa488 probe and thus we obtained FRET-labeled cardiac HMM. FRET between the labeled RLC and the Cy3-ATP reports on the orientation of the myosin light-chain binding domain [19] (Fig. 1A, bound by the ELC and RLC). While binding of the Cy3-ATP stabilizes a pre-powerstroke structural state of the light-chain binding domain, actin stabilizes the post-powerstroke state during the phosphate release phase and prior to the dissociation of the hydrolyzed Cy3-ADP. Cy3-ATP stabilization of the pre-powerstroke state is reflected in a decrease in the time-resolved fluorescence lifetime of the AlexaFluor-488 donor probe and the powerstroke is reflected in the actin-initiated increase in the time-resolved fluorescence (Fig. S3)[19].

We mixed the Cy3-ATP/Alexa488-HMM complex with increasing concentrations of actin containing excess 2 mM MgATP by stopped-flow (Fig. 6A) and acquired time-resolved fluorescence waveforms every 1.0 millisecond during the resulting actin-induced single ATP turnover transient (Fig. 6B). We analyzed the resulting changes in the time-resolved fluorescence decay using a two-distance, structure-based model identical to our prior study [19]. From this we obtain the structural kinetics of the lever-arm rotation in response to actin-activation (Fig. 6C, Table S1).

**Fig. 6.**
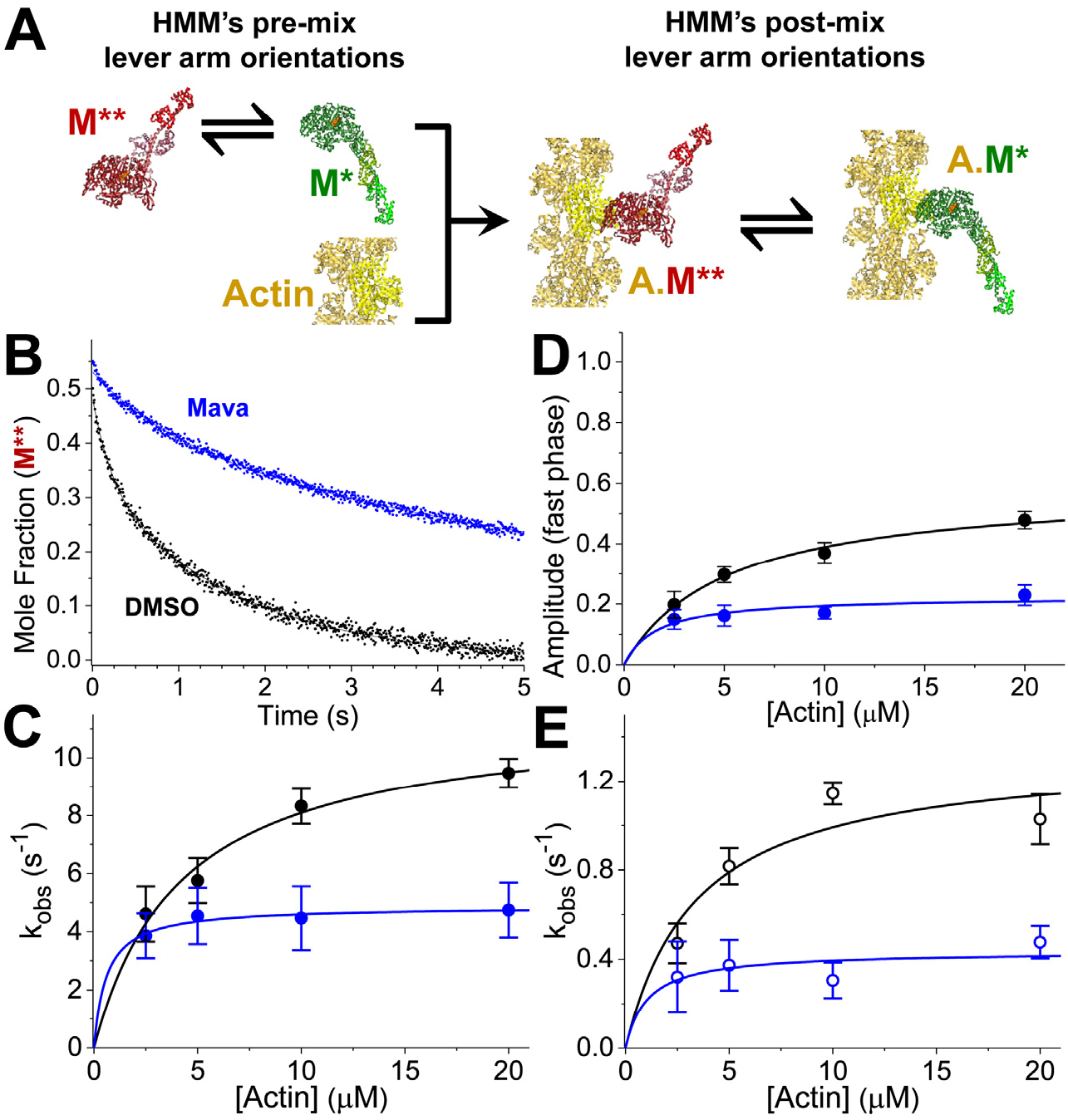
Mavacamten inhibits lever arm rotation in HMM during actin-activation, detected with (TR)^2^FRET. (A) Stopped-flow mix of fluorescently-labeled myosin with actin, to detect FRET between the lever arm and catalytic domain of cardiac HMM during the actin-activated powerstroke. An equilibrium of structural states for myosin (green, red) are depicted before and after stopped-flow mixing. Fluorophores are located on the RLC and nucleotide. S1 shown for simplicity, HMM used. (B) Mole fraction of the M** pre-powerstroke structural state (red), detected with (TR)^2^FRET. Structural transients are fit to two exponentials. (C) Observed rate constants for the fast phase of the actin-activated powerstroke over a range of [Actin]. k_obs,fast,DMSO_ = 11.5s^−1^[Actin]/(4.2μM+[Actin]), k_obs,fast,Mava_ = 4.9s^−1^[Actin]/ (0.6μM+[Actin]). (D) Amplitude of the fast phase: A_fast,DMSO_ = 0.59[Actin]/(5.1μM+[Actin]), A_fast,Mava_ = 0.23[Actin]/ (1.6μM+[Actin]). (E) Slow phase: k_obs.slow,DMSO_ = 1.3s^−1^[Actin]/(3.4μM+[Actin]), k_obs,slow,Myk461_ = 0.4s^−1^[Actin]/(1.0μM+[Actin]). Replicates of n = 6 biochemically independent mixes, each averaging 6-10 shots, two separate preparations of cardiac HMM, error bars represent ± SEM.

In the absence of mavacamten, actin drives lever-arm rotation and the resulting mole-fraction transients for each structural state (pre-powerstroke short distance or post-powerstroke long distance) obtained from fitting the two-state model are best fit to bi-exponential functions (Fig. S3; Fig. 6C-E). The fast phase reflects lever-arm rotation of myosin heads that are primed for activation (Fig. 6C) while the slow phase reflects the equilibration of other HMM species, less readily activated by actin (Fig. 6E).

The maximum observed rate constant for the fast phase of actin-activated lever-arm rotation is 11.5 ± 0.8 s^−1^, and the maximum observed rate constant for the slow phase of actin-activated lever-arm rotation is 1.3 ± 0.2 s^−1^. These rate constants are significantly faster than the fast and slow rate constants for single ATP turnover in the presence of actin (Fig. 2) indicating that actin binding by the two populations of myosin is followed by FRET-detected lever-arm rotation, followed by nucleotide release. Saturating mavacamten significantly reduced these rate constants to 4.9 ± 0.2 s^−1^ (p < 0.0001) and 0.43 ±0.08 s^−1^ (p < 0.001).

We compared the effect of mavacamten on the observed rate constants for actin-activated lever-arm rotation (Fig. 6) to the rate constants for basal single ATP turnover (Fig. 3). At saturating mavacamten and actin, the fast phase of the powerstroke is 23 ± 3% of the total structural transient and is 16-fold faster than the fast phase of basal ATP turnover in the presence of mavacamten (Fig. 6C, D; Fig. 3E). The slow phase of the powerstroke is 77 ± 3% of the transient and is 12-fold faster than the slow phase of basal ATP turnover in the presence of mavacamten (Fig. 6E; Fig. 3D). Because both rate constants for the structural change induced by actin are faster than basal ATP turnover, we conclude that actin is able to bind the mavacamten-inhibited HMM, triggering lever-arm rotation in the fast and slow basal turnover populations. Thus, actin interaction triggers the disruption of the auto-inhibited state of cardiac HMM that mavacamten otherwise stabilizes (Fig. 3).

### Several steps of myosin ATPase cycle are not affected by mavacamten

Additional transient stopped-flow kinetics experiments utilizing (TR)^2^FRET showed that the ATP-induced recovery stroke is not substantially altered by mavacamten (Fig. S4), nor is ATP-binding to myosin or actomyosin, ADP release from actomyosin, myosin dissociation from actomyosin, or the apparent equilibrium constant for ATP hydrolysis measured by acid-quench (Fig. S5, Table S1). Mavacamten’s effects on other steps in myosin’s ATPase cycle, such as phosphate release and actin association in the ADP state, have been reported previously[4].

## Discussion

Regulation of the cardiac myosin thick filament—independent of the actomyosin interaction—is hypothesized to be a critical determinant of contraction and contribute to the Frank-Starling relationship, a fundamental regulator of cardiac performance[26], and to cardiac dysfunction in inherited cardiomyopathies [18, 27]. The structural state of cardiac thick-filaments is hypothesized to be controlled by an evolutionarily-conserved mechanism where the heads of each myosin dimer fold back and interact with their S2 coiled-coil domain and with the thick-filament backbone[13]. This structural state, with myosin heads arranged on the thick filament, has been observed in relaxed muscle with X-ray diffraction[23] and with fluorescence polarization in a specific, head-head conformation termed the interacting heads motif (IHM)[22, 28]. The IHM has been directly observed by electron microscopy studies in isolated filaments[29, 30], and in individual myosin molecules across a variety of myosin II family members[13], including dephosphorylated smooth muscle myosin, and blebbistatin-bound and glutaraldehyde cross-linked skeletal and cardiac myosin[11].

Biochemical studies of permeabilized cardiac and skeletal muscle revealed a population of myosin ATPase sites with dramatically inhibited ATP turnover kinetics compared to isolated myosin S1 analyzed *in vitro* [6, 7, 31]. This population is termed the super-relaxed state (SRX). The SRX biochemical state, defined by very slow single ATP turnover kinetics in the absence of load, is hypothesized to result from the formation of the IHM structural state observed by EM[32]. Neither the SRX nor the IHM have been previously observed in purified cardiac or skeletal myosin in solution without cross-linking or small molecule-based stabilization, and thus the biochemical and structural determinants that are responsible for them have been difficult to study. Our results show that an SRX-like state forms in solution at low ionic strength in the presence of ATP, and that this state controls the kinetics of ATP turnover and actin-activation in two-headed cardiac myosin HMM, summarized in Fig. S1.

Our conclusions are based on measured differences between the biochemical kinetics of single-headed cardiac myosin S1 and two-headed HMM. The steady-state ATPase experiments in Fig. 1 and the transient, actin-activated single ATP turnover studies in Fig. 2 demonstrate that key biochemical differences exist between cardiac S1 and HMM. These differences are indicative of head-head mediated auto-inhibition seen in highly regulated two-headed smooth muscle myosins[33, 34]. These differences also show that mavacamten inhibits both S1 and HMM, but that the compound’s effects on the two-headed HMM protein are distinct and more potent than its effects on S1. Specifically, in HMM, mavacamten stabilizes the slow phase of ATP turnover in the presence (Fig. 2C) and absence of actin (Fig. 3A). It also slows actin-independent ADP release to a greater extent in HMM, indicating that interactions between the two-heads are able to limit ADP dissociation kinetics and that mavacamten stabilizes these interactions (Fig. 3F-H). Furthermore, the compound’s stronger inhibition of HMM compared to S1, indicated by its EC_50_ for steady-state ATPase cycling and actin-activated single ATP turnover (Table S1), shows that the unique structure of the two-headed fragment is preferentially stabilized by the compound.

The temperature and ionic strength dependence studies in Fig. 4 and Fig. 5 support the conclusion that the rate limiting kinetics for ATP turnover in the absence of actin are distinct in HMM compared to S1. The disruption by increasing ionic strength and stability at higher temperatures are consistent with HMM’s two heads interacting in the IHM. These contacts involve putative ionic interactions [18] as well as potential hydrophobic interactions—consistent with stabilization at higher temperature—which have not previously been investigated. Importantly, mavacamten also stabilizes the auto-inhibited state of two-headed HMM in the presence of physiological ionic strengths (Fig. 5D,H).

We directly measured actin-activated structural kinetics in fluorescently-labeled cardiac HMM, detecting movement of the lever arm domain by transient time-resolved FRET (Fig. 6, Fig. S3). The slow phase of actin-activated lever arm rotation detected in this experiment is significantly faster than mavacamten-saturated HMM’s basal ATP turnover kinetics (Fig. 3A, C-E). This result indicates that at least one head of the auto-inhibited HMM is still able to interact with actin. It also indicates that actin interaction with the auto-inhibited HMM accelerates the structural disruption of the inhibited state. This provides an explanation for why the IHM has been difficult to observe in cardiac and skeletal myosin by EM—interactions with the UV-treated carbon surface or charged mica surface could disrupt it in the absence of cross-linking[13] or stabilizing small-molecules[11].

Based on our results, we propose a mechanism for actin-activation of two-headed myosin in the presence and absence of mavacamten (Fig. 7). In this mechanism auto-inhibited myosin heads transition from a folded IHM state that is docked on the thick-filament backbone (state 1) to an IHM state with S2 extended off the filament backbone (state 2). The IHM heads then “open” (state 3) and become available for actin-activation (state 4). Myosin in the IHM can thus be activated by following the pathway (1) ↔ (2) ↔ (3) ↔ (4) (Fig. 7). Alternatively, actin interaction accelerates IHM head opening if one of the heads—we hypothesize the free IHM head[13] (Fig. 1A)—weakly interacts with available myosin binding sites on the actin thin-filament (state 5 in Fig. 7). Thus, activation can also proceed via the pathway (1) ↔ (2) ↔ (5) ↔ (4). This second, actin-activated pathway is supported by our biochemical and structural kinetics data and also depicted in the myosin ATPase cycle in Fig. S1. The “SRX – Heads up” state (2) is likely present in muscle fibers because we have observed it in tissue-purified HMM, and it is stable at physiological temperatures (Fig. 4). This state is weakened at physiological ionic strengths (Fig. 5), indicating that additional protein-protein interactions are likely required for its full stabilization in the cardiac sarcomere. Similar arguments were made in a previous paper based on head-to-S2 binding experiments performed by microscale thermophoresis[18]. We find that mavacamten shifts the apparent equilibrium towards state (2) in the presence of ATP as well as ADP (Fig. 3A,E). Mavacamten inhibits the kinetics of the IHM opening transition given that it significantly decreases the rate constant for the slow phase of HMM basal single ATP turnover more than the slow phase of S1 (Fig. 3I) (p < 0.0001).

**Fig. 7.**
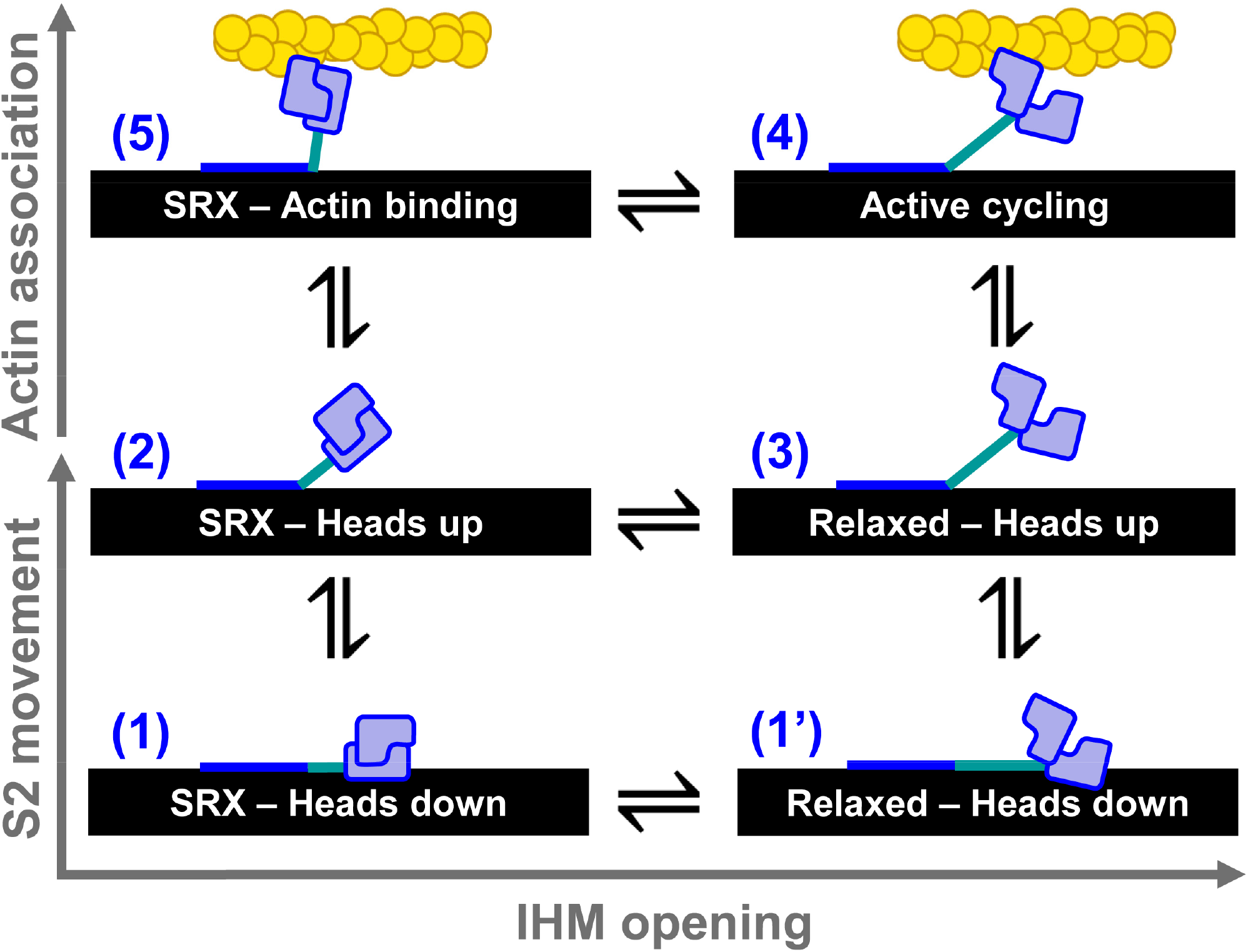
Structural model for thick filament regulation. (1) Electron microscopy data supports an interacting-heads motif folded back against the thick filament (black). (1’) The heads may open while the S2 is on the thick filament. (2) Hypothesized state based on our detection of an SRX-like state in HMM. (3) Relaxed myosin with heads splayed and ready to interact with actin. (4) Active actomyosin cycling. (5) Hypothesized state based our detection of actin accelerating the auto-inhibited basal ATP turnover of HMM, and thus likely unfolding the IHM.

Under concentrations of mavacamten sufficient to fully inhibit ATPase cycling, we propose that HMM still follows the pathway through states (1) ↔ (2) ↔ (5) ↔ (4), because even though mavacamten stabilizes the auto-inhibited state, actin is still able to accelerate ATP turnover. This conclusion is based on the observation that the slowest phases of lever-arm rotation detected by (TR)^2^FRET (0.43 s^−1^), and actin-activated single ATP turnover (0.04 s^−1^), are both substantially faster than the fast phase (p < 0.0001, p < 0.0002) or slow phase (p < 0.0001, p < 0.0001) of mavacamten inhibited basal single ATP turnover by cardiac HMM.

## Conclusion

We have detected a biochemically auto-inhibited state in tissue-purified cardiac HMM. The kinetics and energetics of this state are consistent with the SRX observed in permeablized myocardium and thus we believe we have observed the SRX state in a striated muscle myosin in solution for the first time. Importantly, mavacamten stabilizes this auto-inhibited state, revealing a new mechanism for a drug’s inhibition of two-headed cardiac myosin. Actin interaction is able to disrupt the auto-inhibited state and accelerate the rate-limiting structural transition of head-head splaying that limits ATP turnover in HMM, unlike in S1. Thus, we hypothesize that weak interaction with actin disrupts the SRX-like, auto-inhibited state of cardiac myosin in the myocardium as well. We also propose that because weak-actin interactions disrupt the auto-inhibited state in solution, factors which control extension of the IHM state off the filament backbone (states (1) ↔ (2), Fig. 7) will be important in contracting cardiac muscle. Mechanisms likely regulating this transition include RLC phosphorylation[35], myosin-binding protein-C[31], thick-filament mechano-sensing[36], and small molecules[37].

## Methods

### Steady-state ATPase activity

The actin-activated MgATPase activity of purified cardiac myosin HMM or S1 was measured using an NADH-coupled assay [20] performed at 25 ºC in 10 mM Tris pH 7.5, 2 mM MgCl_2_ with 1.0 mM DTT. The reaction mix contained 0.2 μM HMM or 0.4 μM S1, varied [actin], and 0.2 mM NADH, 0.5 mM PEP, 2.1 mM ATP, 10 U/mL LDH, 40 U/mL PK, HMM (200 nM). We acquired absorbance at 340 nm every 10 seconds for 120 seconds total using a Beckman-Coulter DU640B spectrophotometer.

### Transient kinetics

Transient biochemical experiments with steady-state fluorescence detection (total fluorescence intensity) were performed on an *Applied Photophysics* stopped-flow spectrophotometer capable of single-mix and sequential-mix experiments with water bath temperature control. All experiments performed at 25 °C unless otherwise stated. The single-mix dead time for this instrument is 1.3 ms. All buffers were filtered and then degassed for 30 minutes under high-vacuum prior to use.

Single ATP turnover experiments with mant-ATP were performed with both sequential and single stopped flow mixes. Samples were excited at 280 nm with a Xe lamp and monochromator, and detected through a 400 nm long-pass filter.

### Statistics and error analysis

Individual, representative traces shown throughout the manuscript depict the average of 6-10 shots of the stopped-flow. All experiments were performed in replicates of n = 4 to 9, and as biochemically independent experiments. Three separate preparations of cardiac HMM from separate bovine myocardia obtained unfrozen from Pel-Freez Biologicals contributed to the experiments throughout this manuscript; error bars represent ± SEM. The student’s t-test was used to determine statistical significance of measured parameters, tabulated in Table S1.

A complete discussion of all methods, including the extraction, digestion, purification and expression of the proteins utilized in this study, the steady-state and transient kinetics methodologies, and (TR)^2^FRET data acquisition and analysis are included in the SI.

## ACKNOWLEDGEMENTS

This study was supported by grants to D.D.T from NIH (R01AR32961, R42DA037622) and to J.M.M from the American Heart Association (14SDG20480032). J.A.R. was supported by a Graduate Excellence Fellowship from the University of Minnesota. We thank John Lipscomb for use of his sequential stopped-flow instrument. We thank Piyali Guhathakurta for assistance purifying cardiac myosin and data acquired on S1 ATPase activity in Figure 1C, Osha Roopnarine for experimental assistance acquiring pyrene-actin quenching by S1 data in Figure 1E, and Sami Chu for assistance with molecular biology and purification of bovine RLC. We also thank Christopher Yengo, Steven Rosenfeld, Lien Phung, Sami Chu, Yahor Savich, Roger Cooke, James Spudich, Kathy Ruppel, Darshan Trivedi, Suman Nag, Linda Song, and Chao Liu for generous helpful discussions during key phases of this work.

## Supplemental Methods

### Protein purification and labeling

#### Myosin

Bovine cardiac myosin was purified following procedures based on Margossian et al. [38]. Cow hearts were obtained on wet ice from *Pel-Freez*. The entire purification process was performed in a 4 ºC cold room. A thorough description is given in our previous publications [19].

#### HMM digestion

Myosin prepared as described in Rohde et al. [19] was thawed and brought to 2 mM MgCl_2_ in its freezing buffer before digestion to HMM with α-chymotrypsin (*Sigma-Aldrich*, 0.025 mg/ml final concentration) for 10 minutes at 25 ºC, followed by addition of pefabloc (*Roche*, 5 mM final concentration) and then dialyzed into 10 mM Tris pH 7.5 with 2 mM MgCl_2_ and then purified by Q-sephadex ion-exchange chromatography. The column was equilibrated in 10 mM Tris pH 8.0 at 4.0 C, the digested myosin loaded and then eluted in a gradient of 0-300 mM KCl, 10 mM Tris pH 8.0, over 300 mL at 1.5 mL/min, collecting 4 mL fractions (AKTA Prime Plus, GE). Fractions were evaluated by SDS-Page and those containing intact HMM without contaminants, pooled for experiments. The pooled HMM was dialyzed into buffers utilized for each experiment.

#### RLC

Expression, labeling and storage of recombinant Bovine ventricular regulatory light chain with a single engineered cysteine at position 105, is described in our previous work [19].

#### Actin

Actin was purified from rabbit skeletal muscle by acetone dehydration followed by extraction into ice cold water as described in our previous work [39] and then polymerized in 10 mM Tris pH 7.5, 2 mM MgCl_2_, 0.5 mM ATP and stored on ice prior to use. Prior to use, the F-actin was stabilized with a 1:1.3 stochiometric excess of phalloidin (*Sigma Aldrich*), followed by 48 h dialysis (3 buffer changes) into 10 mM Tris pH 7.0 with 2 mM MgCl_2_.

#### Protein and dye concentration

The Bradford protein concentration assay utilizing a known BSA protein standard was used throughout this study to determine protein concentrations. Reagents for this assay were purchased from *Biorad*. The extinction coefficient for the AF488 dye is 73,000 at 495 nm, and for Cy3 is 136,000 at 570 nm, per the manufacturer’s specifications.

#### Exchange

We exchanged the AF488-labeled RLC onto HMM by combining the two proteins (4.5 molar excess RLC to HMM) in 50 mM Tris pH 7.5, 120 mM KCl, 2 mM DTT, 12 mM EDTA [40] and then incubated the reaction mix for 30 minutes at 30°C. After the incubation, we adjusted the reaction to 12 mM MgCl_2_ and then incubated the mixture on ice for 15 minutes followed by dialysis into 10 mM Tris pH 7.0, 30 mM KCl, 2 mM MgCl_2_ prior to gel filtration to remove free RLC.

#### Buffers and solutions

All experiments, unless otherwise noted, were performed in 10 mM Tris pH 7.5, 2 mM MgCl_2_ at 25 ºC.

#### Chemicals

Mavacamten (Myk-461) was custom synthesized by EAG Laboratories; purity was >97% by proton nuclear magnetic resonance (^1^H NMR) and 99.36% by liquid chromatography/mass spectrometery (LC/MS). ATP (Adenosine 5’-triphosphate disodium salt hydrate, Grade 1 >99 %) and ADP (Adenosine 5’-diphosphate sodium >95 %) were purchased from *Sigma Aldrich*. Mant-ATP (2’-(or-3’)-O-(N-methylanthraniloyl) adenosine 5’-triphosphate, trisodium salt) was purchased from *Thermo Fisher Scientific*. All other chemicals where purchased from *Sigma Aldrich*.

#### Steady-state ATPase activity

We measured the actin-activated MgATPase activity of the purified cardiac myosin HMM using an NADH-coupled assay [20] performed at 25 ºC in 10 mM Tris pH 7.5, 2 mM MgCl_2_. The reaction mix contained varied [actin], and 0.2 mM NADH, 0.5 mM PEP, 2.1 mM ATP, 10 U/mL LDH, 40 U/mL PK, HMM (200 nM). We acquired absorbance at 340 nm every 10 seconds for 120 seconds total using a Beckman-Coulter DU640B spectrophotometer.

#### Transient kinetics

Transient biochemical experiments with steady-state fluorescence (total fluorescence intensity) detection were performed on an *Applied Photophysics* stopped-flow spectrophotometer capable of sequential mixing experiments with water bath temperature control. All experiments performed at 25 °C unless otherwise stated. The single-mix dead time for this instrument is 1.3 ms, calibrated using fluorescence enhancement of 8-hydroxyquinoline following Mg^+2^ binding under pseudo first-order kinetics conditions [41]. All buffers were filtered and then degassed for 30 minutes under high-vacuum prior to use.

Transient time-resolved FRET (millisecond-resolved transient biochemical experiments with nanosecond-resolved fluorescence detection), (TR)^2^FRET, was measured using a transient time-resolved fluorescence spectrophotometer [42–44]. This instrument utilizes a *Biologic USA* SFM/20 single-mix stopped-flow accessory coupled to our transient time-resolved fluorescence spectrophotometer. The dead time for the instrument was 1.8 ms, calibrated using the 8-hydroxyquinoline + Mg^+2^ control reaction [41]. For experiments mixing equilibrated myosin in the presence of 10 molar excess ATP with actin containing 1 mM MgATP, we loaded the actin into syringe A, followed by a freshly prepared 600 μL mixture of myosin + Cy3-ATP in syringe B and then immediately mixed with the actin in syringe A.

Single-turnover experiments with mant-ATP were performed with a sequential stopped flow. For mant-ATP: excitation at 280 nm and detection through a 400 nm long-pass filter.

Pyreneiodoacetimide (PIA) labeled actin was prepared as described in [45]. Pyrene-actomyosin association in the absence of nucleotide, pyrene-actomyosin dissociation in the presence of 50-molar excess of unlabeled actin, and ATP-induced dissociation of actomyosin was performed as described in [20].

#### Hydrolysis of ATP

We detected free phosphate in solution as described in [46]. 12.5 μM cardiac HMM (25 μM heads) was manually mixed with 20 μM ATP (Sigma-Aldrich), allowed to incubate for 5.0 seconds and quenched with 0.6 M perchloric acid, and detected with malachite green as described in [47]. Experiments were performed at room temperature, 22-23 °C.

#### Time-resolved fluorescence resonance energy transfer (TR-FRET) and transient TR-FRET, (TR)^2^FRET

Fit parameters of the two-distance model are given in Table S1 and a detailed description of data fitting is described in Rohde et al. [19].

#### (TR)^2^FRET

The TRF and (TR)^2^F spectrometers, originally described in our previous work [42–44], transiently digitize the time-resolved fluorescence emission following a 1 ns laser pulse. The laser used in this study is an artisanal 473 nm microchip laser (FP2-473-3-5) with an LD-702 controller hand crafted by *Concepts Research Corporation*, in WI, operating at 5 KHz repetition frequency. Thus samples are excited every 0.2 ms. For equilibrium and steady-state biochemical conditions, 1000 replicate waveforms were signal-averaged prior to analysis. For transient time-resolved measurements acquired after rapid mixing by stopped-flow, 5 waveforms were averaged every 1 ms. Total time-resolved fluorescence was measured with the emission polarizer set to the magic angle (54.7°) or removed.

### (TR)^2^FRET Data Analysis

#### Total fluorescence

We determined the total fluorescence emission for FRET samples by integrating the (TR)^2^FRET waveforms over the nanosecond decay time after subtracting the pre-trigger dark current, ∼5% in amplitude compared to the maximum waveform intensity.

#### TR-FRET

TRF waveforms from donor and FRET-labeled samples were analyzed as described in our previous publications [42–44] Eq. 1–13, paraphrased below. The measured time-resolved fluorescence waveform, *I*(t) (Eq 1),

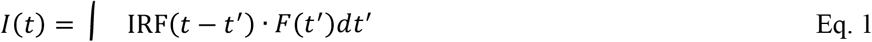

is a function of the nanosecond decay time, t, and is modeled as the convolution integral of the measured instrument response function, IRF(t), and the fluorescence decay model, *F*(t). The fluorescence decay model (Eq. 2)

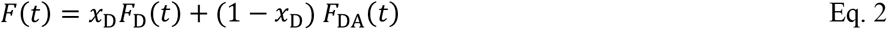

is a linear combination of a donor-only fluorescence decay function, *F*_D_(t) and an energy transfer-affected donor fluorescence decay, *F*_DA_(t). The donor decay *F*_D_(t) is a sum of exponentials (Eq. 3)

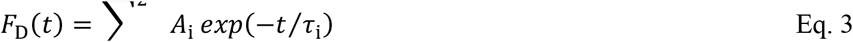

with discrete lifetime species τ_i_ and pre-exponential mole fractions *A*_i_. For the Alexa-488 donor two exponentials were required to fit the observed fluorescence. The energy transfer-affected donor decay function, *F*_DA_(t) (Eq. 4),

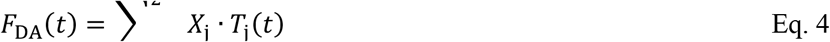

is a sum over multiple structural states (*j*) with mole fractions *X*_j_, represented by FRET-affected donor fluorescence decays *T*_j_(t). The increase in the donor decay rate (inverse donor lifetime) due to FRET is given by the Förster equation

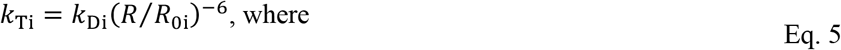

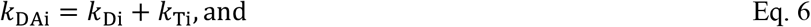

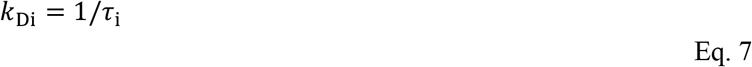

We modeled TR-FRET assuming that each structural state *j* (Eq. 4) corresponds to a Gaussian distribution of interprobe distances, ρ_j_(*R*):

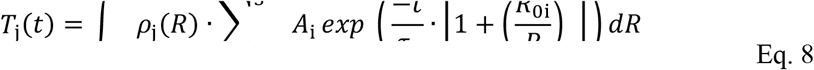

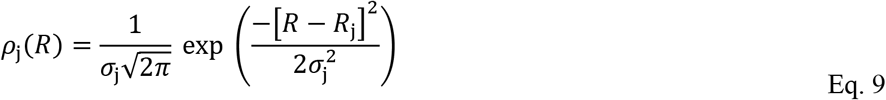

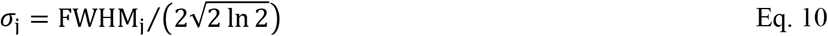

As with our previous work [42–44], *R*_0i_ is calculated according to Eq. 11 from the spectral overlap integral, *J*, the orientation-sensitive term *κ*^2^, the refractive index n, and the donor quantum yield *Q*_Di_ (Eq. 12–14). 〈*Q*_D_〉 was measured as 0.91 ± 0.01, by comparison to a quinine sulfate fluorescence standard in 50 mM H_2_SO_4_ at 25°C according to Eq. 14 (1, 4).

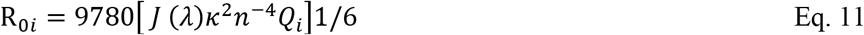

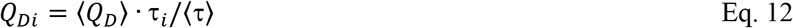

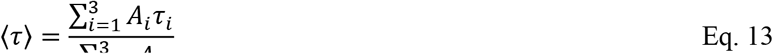

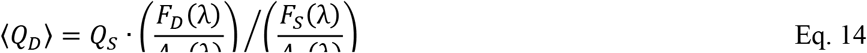

Together, the donor fluorescence (*A*_i_, τ_i_) and distance terms (*R*_*j*_, σ_j_) in our analysis were shared globally between all waveforms containing FRET-labeled samples. *R*_j_ and σ_j_ were allowed to vary between 0.5 nm and 15.0 nm. The average Alexa-488/CY3 *R*_0_, (6.7 nm in this study) was determined according to Eq. 11–14. The distance-dependent terms *R*_j_ (Eq.9) and σ_j_ (Eq. 10) define unique structural states of the LCD. The mole fraction terms *X*_j_ were allowed to vary independently in each waveform. Thus, changes in *X*_j_ reflect changes in the relative populations of the structural states (*j*) as the biochemical state is varied under equilibrium, steady-state, or transient conditions.

We determined the number of donor lifetimes (*i*) and structural states (*j*) that are present in each sample by fitting a set of models with the number of donor lifetime states, *i* increasing from 1 to 4, and the number of structural states, *j*, increasing from 1 to 4. For each model we test a distribution of energy transfer rates, with σ_*j*_ allowed to vary, as well as discrete energy transfer rates where σ → 0. The final model (*i*_max_ *= 2, j*_max_ *= 2, σ > 0*) was determined by evaluating the dependence of the minimized *χ*^2^ on the number of free parameters in the global model and by the resolution of the *χ*^2^ error surface support plane with a confidence intervals of 67%.

---

**Table S1.** Steady-state and transient kinetics measured in this paper. HMM and DMSO (black), S1 and DMSO (red), HMM and 10 μM Mava (blue), S1 and 10 μM Mava (purple). Student’s T-Test reporting: N.S. for p > 0.05, **\*** for p ≤ 0.05 ≥ p, **\*\*** for p ≤ 0.01, **\*\*\*** for p ≤ 0.001.

| **Table S1.** Steady-state and transient kinetics measured in this paper. HMM and DMSO (black), S1 and DMSO (red), HMM and 10 μM Mava (blue), S1 and 10 μM Mava (purple).<br>Student's T-Test reporting: N.S. for $p > 0.05$, * for $p \leq 0.05 \geq p$, ** for $p \leq 0.01$, *** for $p \leq 0.001$. | | | |
| --- | --- | --- | --- |
| Experiment, Parameters | Myosin fragment, ± DMSO or Mava | | Stat. signif. |
| **HMM steady-state, actin-activated NADH-coupled ATPase** | **HMM, DMSO** | **HMM, Mava** |  |
| $k_{cat}$ ($s^{-1}$), $n=6$, Error of 3 fit hyperbolas. | $2.02 \pm 0.12$ | $0.48 \pm 0.04$ | *** |
| $K_{0.5,Actin}$ (μM) | $17.6 \pm 2.8$ | $35.3 \pm 6.3$ | * |
| Mavacamten $EC_{50}$ (μM), $n=4$ | N/A | $0.23 \pm 0.02$ | |
| **S1 steady-state, actin-activated NADH-coupled ATPase** | **S1, DMSO** | **S1, Mava** |  |
| $k_{cat}$ ($s^{-1}$), $n=6$, Error of 3 fit hyperbolas. | $3.64 \pm 0.41$ | $1.35 \pm 0.08$ | *** |
| $K_{0.5,Actin}$ (μM) | $24.4 \pm 7.4$ | $20.6 \pm 8.1$ | N.S. |
| Mavacamten $EC_{50}$ (μM) | N/A | $0.47 \pm 0.006$ [4] | |
| **HMM steady-state, basal NADH-coupled ATPase** | **HMM, DMSO** | **HMM, Mava** |  |
| Activity ($s^{-1}$), Repli.: $n=3$, Error: ± SEM. | $0.016 \pm 0.005$ | $0.003 \pm 0.001$ | * |
| **S1 steady-state, basal NADH-coupled ATPase** | **S1, DMSO** | **S1, Mava** |  |
| Activity ($s^{-1}$), Repli.: $n=3$, Error: ± SEM. | $0.028 \pm 0.007$ | $0.006 \pm 0.002$ | ** |
| **HMM steady-state pyrene-actin quenching** | **Acto-HMM** | **Acto-S1** |  |
| Normalized fluorescence (A.U.), $n=4$, Error: ± SEM. | $0.29 \pm 0.03$ | $0.34 \pm 0.02$ | N.S. |
| **S1 steady-state pyrene-actin quenching** | **ActoHMM+ATP** | **ActoS1 + ATP** |  |
| Normalized fluorescence (A.U.), $n=4$, Error: ± SEM. | $0.89 \pm 0.02$ | $0.95 \pm 0.01$ | * |
| **HMM actin-activated single-turnover of mant-ATP** | **HMM, DMSO** | **HMM, Mava** |  |
| $k_{fast}$ ($s^{-1}$), $n=6$, Error of 3 hyperbolic fits. | $3.80 \pm 0.33$ | $0.30 \pm 0.03$ | *** |
| $K_{0.5,Actin}$ (μM) | $5.3 \pm 1.7$ | $2.3 \pm 0.5$ | N.S. |
| $k_{slow}$ ($s^{-1}$) | $0.55 \pm 0.03$ | $0.037 \pm 0.007$ | *** |
| $K_{0.5,Actin}$ (μM) | $5.4 \pm 1.1$ | $4.7 \pm 1.5$ | N.S. |
| $A_{fast}$, fraction (A.U.) | $0.77 \pm 0.03$ | $0.48 \pm 0.04$ | ** |
| Mavacamten $EC_{50}$ (μM) | N/A | $0.36 \pm 0.08$ | |
| **S1 actin-activated single-turnover of mant-ATP** | **S1, DMSO** | **S1, Mava** |  |
| $k_{fast}$ ($s^{-1}$), $n=6$, Error of 3 hyperbolic fit. | $5.97 \pm 0.80$ | $0.80 \pm 0.11$ | ** |
| $K_{0.5,Actin}$ (μM) | $16.1 \pm 5.0$ | $12.6 \pm 4.2$ | N.S. |
| $k_{slow}$ ($s^{-1}$) | $0.28 \pm 0.04$ | $0.089 \pm 0.002$ | ** |
| $K_{0.5,Actin}$ (μM) | $2.7 \pm 0.9$ | $1.3 \pm 0.3$ | N.S. |
| $A_{fast}$, fraction (A.U.) | $0.87 \pm 0.02$ | $0.83 \pm 0.01$ | N.S. |
| **HMM basal single-turnover of mant-ATP** | **HMM, DMSO** | **HMM, Mava** |  |
| $k_{fast}$ ($s^{-1}$), $n=9$, Error: ± SEM. | $0.021 \pm 0.003$ | $0.0076 \pm 0.0012$ | *** |
| $k_{slow}$ ($s^{-1}$) | $0.0053 \pm 0.0010$ | $0.0008 \pm 0.0002$ | *** |
| $A_{slow}$, fraction (A.U.) | $0.55 \pm 0.04$ | $0.88 \pm 0.01$ | *** |
| **S1 basal single-turnover of mant-ATP** | **S1, DMSO** | **S1, Mava** |  |
| $k_{fast}$ ($s^{-1}$), $n=9$, Error: ± SEM. | $0.032 \pm 0.002$ | $0.0038 \pm 0.0012$ | *** |
| $k_{slow}$ ($s^{-1}$) | $0.0075 \pm 0.0010$ | $0.0013 \pm 0.0003$ | *** |
| $A_{slow}$, fraction (A.U.) | $0.17 \pm 0.04$ | $0.32 \pm 0.03$ | ** |
| **HMM basal single-turnover of ATP, chased with mant-ATP** | **HMM, DMSO** | **S1, DMSO** |  |
| $k_{fast}$ ($s^{-1}$), $n=6$, Error: ± SEM. | $0.028 \pm 0.005$ | $0.025 \pm 0.002$ | N.S. |
| $k_{slow}$ ($s^{-1}$) | $0.0039 \pm 0.0011$ | $0.0045 \pm 0.0006$ | N.S. |
| $A_{slow}$, fraction (A.U.) | $0.81 \pm 0.02$ | $0.27 \pm 0.03$ | *** |
| **AF488-RLC-HMM basal single-turnover of mant-ATP** | **HMM** | **AF488-HMM** |  |
| $k_{fast}$ ($s^{-1}$), $n=9$ for HMM, $n=4$ for AF488-HMM, Error: ± SEM. | $0.021 \pm 0.003$ | $0.011 \pm 0.002$ | N.S. |
| $k_{slow}$ ($s^{-1}$) | $0.0053 \pm 0.0010$ | $0.0037 \pm 0.0007$ | N.S. |
| $A_{slow}$, fraction (A.U.) | $0.55 \pm 0.04$ | $0.47 \pm 0.03$ | N.S. |
| **HMM basal mant-ADP release** | **HMM, DMSO** | **HMM, Mava** |  |
| $k_{fast}$ ($s^{-1}$), $n=9$, Error: ± SEM. | $0.45 \pm 0.01$ | $0.16 \pm 0.01$ | *** |
| $k_{slow}$ ($s^{-1}$) | $0.12 \pm 0.01$ | $0.025 \pm 0.003$ | *** |
| Amplitude<sub>slow</sub>, fraction (A.U.) | $0.37 \pm 0.02$ | $0.62 \pm 0.02$ | *** |
| **S1 basal mant-ADP release** | **S1, DMSO** | **S1, Mava** |  |
| $k_{fast}$ ($s^{-1}$), $n=9$, Error: ± SEM. | $0.41 \pm 0.02$ | $0.20 \pm 0.01$ | *** |
| $k_{slow}$ ($s^{-1}$) | $0.08 \pm 0.01$ | $0.04 \pm 0.01$ | * |
| Amplitude<sub>slow</sub>, fraction (A.U.) | $0.30 \pm 0.03$ | $0.28 \pm 0.01$ | N.S. |
| **Basal mant-ATP turnover, temperature dependence** | **HMM** | **S1** |  |
| **Eyring equation:** $k = T \cdot e^{\left(\frac{-\Delta H^\ddagger}{RT} + b\right)}$, $R = 8.314(10^{-3}) \text{ kJ} \cdot \text{K}^{-1} \cdot \text{mol}^{-1}$ | | | |
| Fast phase: $\Delta H^\ddagger$ (kJ/mol), $n=4$, Error of fit. | $35.3 \pm 1.5$ | $30.9 \pm 3.3$ | N.S. |
| Fast phase: $b$, constants and $\Delta S^\ddagger$ (dimensionless) | $5.3 \pm 0.6$ | $3.6 \pm 1.4$ | N.S. |
| Slow phase: $\Delta H^\ddagger$ (kJ/mol) | $15.9 \pm 1.3$ | $15.1 \pm 8.9$ | N.S. |
| Slow phase: $b$, constants and $\Delta S^\ddagger$ (dimensionless) | $-4.3 \pm 0.5$ | $-3.9 \pm 3.7$ | N.S. |
| Amplitude<sub>slow</sub>, fraction at 35 °C (A.U.), $n=4$, Error: ± SEM. | $0.51 \pm 0.03$ | $0.07 \pm 0.03$ | *** |
| **Basal mant-ATP turnover, KCl dependence** | **HMM** | **S1** |  |
| Average $k_{fast}$ ($s^{-1}$), $n=4$, Error: ± SEM. | $0.022 \pm 0.001$ | $0.031 \pm 0.002$ | ** |
| Average $k_{slow}$ ($s^{-1}$) | $0.0062 \pm 0.0004$ | $0.0069 \pm 0.0013$ | N.S. |
| Amplitude<sub>slow</sub>, fraction at 100 mM KCl (A.U.), Error of fit. | $0.23 \pm 0.10$ | $0.24 \pm 0.09$ | N.S. |
| **Basal mant-ATP turnover, 100 mM KCl and 10 μM Mava** | **HMM** | **S1** |  |
| $k_{fast}$ ($s^{-1}$), $n=4$, Error: ± SEM. | $0.0037 \pm 0.0008$ | $0.0049 \pm 0.0005$ | N.S. |
| $k_{slow}$ ($s^{-1}$) | $0.0009 \pm 0.0001$ | $0.0011 \pm 0.0002$ | N.S. |
| $A_{slow}$, fraction (A.U.) | $0.59 \pm 0.03$ | $0.46 \pm 0.02$ | * |
| **Powerstroke, (TR)<sup>2</sup>FRET, fluorescently-labeled RLC** | **HMM, DMSO** | **HMM, Mava** |  |
| Actin-induced lever arm rotation ($s^{-1}$), $n=6$, Error of 3 fits. | $11.49 \pm 0.81$ | $4.87 \pm 0.17$ | ** |
| Actin-induced rotation, $K_{0.5, \text{Actin}}$ (μM) | $4.20 \pm 0.89$ | $0.6 \pm 0.2$ | * |
| Slow phase, lever arm rotation ($s^{-1}$) | $1.33 \pm 0.24$ | $0.43 \pm 0.08$ | * |
| Slow phase, $K_{0.5, \text{Actin}}$ (μM) | $3.42 \pm 2.05$ | $1.0 \pm 1.3$ | N.S. |
| Maximum fit Amplitude, fast phase (normalized) | 0.59 ± 0.04 | 0.23 ± 0.03 | ** |
| Maximum fit Amplitude, K<sub>0.5,Actin</sub> (μM) | 5.1 ± 0.8 | 1.6 ± 1.0 | N.S. |
| **Distance Distribution, (TR)<sup>2</sup>FRET** | **HMM, DMSO** | **HMM, Mava** |  |
| M** (Distance 1), *n=28, Error is avg. surface plane error.* | 6.54 ± 0.55 | Globally-fit |  |
| FWHM 1 (nm) | 1.87 ± 0.62 | “ |  |
| M* (Distance 2) (nm) | 10.53 ± 1.15 | “ |  |
| FWHM 2 (nm) | 2.20 ± 0.82 | “ |  |
| **Donor-only fluorescence, AF488, (TR)<sup>2</sup>FRET** | **HMM, DMSO** | **HMM, Mava** |  |
| τ<sub>1</sub> (ns), *n=12, Error is avg. surface plane error.* | 3.791 ± 0.008 | Globally-fit |  |
| A<sub>1</sub> (A.U.) | 0.645 ± 0.029 | “ |  |
| τ<sub>2</sub> (ns) | 0.493 ± 0.011 | “ |  |
| **Pyrene-A.M diss. (50x unlabeled-actin chase)** | **HMM, DMSO** | **HMM, Mava** |  |
| k<sub>1</sub> (s<sup>-1</sup>), *n=6, Error: ± SEM.* | 0.052 ± 0.004 | 0.052 ± 0.006 | N.S. |
| A<sub>1</sub>, fraction (A.U.) | 0.23 ± 0.01 | 0.24 ± 0.01 | N.S. |
| k<sub>2</sub> (s<sup>-1</sup>) | 0.0071 ± 0.0001 | 0.0075 ± 0.0001 | * |
| **Mant-ATP binding Myosin** | **HMM, DMSO** | **HMM, Mava** |  |
| k<sub>1</sub> (s<sup>-1</sup>), *n=3, Error: ± SEM.* | 24.3 ± 0.8 | 29.0 ± 1.0 | * |
| A<sub>1</sub>, fraction (A.U.) | 0.69 ± 0.01 | 0.77 ± 0.02 | * |
| k<sub>2</sub> (s<sup>-1</sup>) | 1.42 ± 0.6 | 1.75 ± 0.9 | N.S. |
| **Mant-ATP binding Actomyosin** | **HMM, DMSO** | **HMM, Mava** |  |
| k<sub>1</sub> (s<sup>-1</sup>), *n=6, Error: ± SEM.* | 309 ± 110 | 143 ± 22 | N.S. |
| A<sub>1</sub>, fraction | 0.27 ± 0.01 | 0.28 ± 0.02 | N.S. |
| k<sub>2</sub> (s<sup>-1</sup>) | 25.6 ± 0.9 | 20.9 ± 0.7 | ** |
| A<sub>2</sub>, fraction | 0.63 ± 0.02 | 0.61 ± 0.01 | N.S. |
| k<sub>3</sub> (s<sup>-1</sup>) | 6.4 ± 2.4 | 1.8 ± 0.3 | N.S. |
| **Mant-ADP release from actoHMM, chased with ATP** | **HMM, DMSO** | **HMM, Mava** |  |
| k<sub>1</sub> (s<sup>-1</sup>), *n=6, Error: ± SEM.* | 66.8 ± 0.8 | 64.5 ± 2.3 | N.S. |
| A<sub>1</sub>, fraction | 0.93 ± 0.01 | 0.92 ± 0.01 | N.S. |
| k<sub>2</sub> (s<sup>-1</sup>) | 2.4 ± 0.4 | 2.4 ± 0.3 | N.S. |
| **Recovery stroke, (TR)<sup>2</sup>FRET** | **HMM, DMSO** | **HMM, Mava** |  |
| Fast phase, ATP-induced lever arm rotation, k<sub>cat</sub> (s<sup>-1</sup>) | 58.5 ± 11.7 | 42.1 ± 4.7 | N.S. |
| ATP-induced lever arm rotation K<sub>0.5</sub> [Cy3ATP] (μM) | 10.4 ± 3.2 | 8.3 ± 1.5 | N.S. |
| Slow phase, k<sub>cat</sub> (s<sup>-1</sup>) | 1.97 ± 0.38 | 1.24 ± 0.20 | N.S. |
| Slow phase K<sub>0.5</sub> [Cy3ATP] (μM) | 1.56 ± 0.93 | 1.29 ± 0.70 | N.S. |
| Mavacamten EC<sub>50</sub> (μM) | N/A | 27.5 ± 4.7 |  |
| **Steady state hydrolysis** | **HMM, DMSO** | **HMM, Mava** |  |
| [P<sub>i</sub>] / [ATP] (no units), *n=6, Error: ± SEM.* | 0.62 ± 0.03 | 0.55 ± 0.01 | N.S. |

**Fig. S1.**
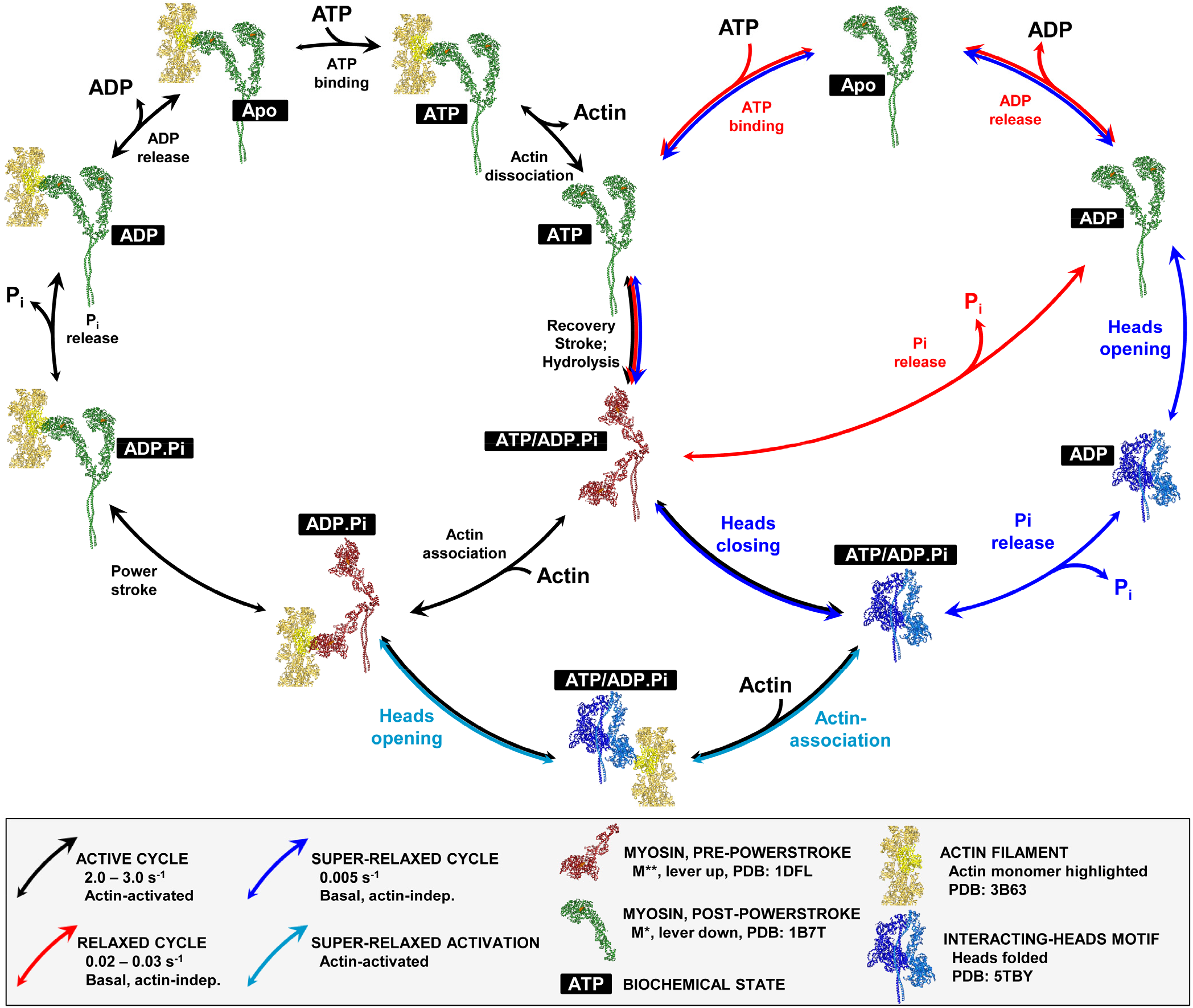
ATP kinetic cycle of the soluble, two-headed cardiac myosin fragment HMM. Proposed kinetic cycle of cardiac HMM, emphasizing actin-activated turnover of ATP (black arrows), relaxed (basal) actin-independent ATP cycling (red arrows), super-relaxed ATP cycling (blue arrows), and actin-accelerated unfolding of the super-relaxed state (light blue arrows). Myosin’s biochemical state is indicated in white letters with black background, with “apo” indicating that no nucleotide is bound. We propose that two-headed myosin’s actin-activated ATP cycle (left loop, black arrows) includes the interacting heads motif (lower center loop, black arrows), resulting in decreased steady-state and transient ATP cycling in HMM compared to S1, consistent with data presented (Fig. 1, Fig. 2). We hypothesize that the free head is able to associate with actin (Fig. 2, Fig. 6). We also propose that P_i_ release precedes the heads opening, followed by ADP release during basal cycling (blue arrows, right edge). This is based on HMM’s and S1’s identical rates of basal ADP release, and in the same transient experiment, mavacamten’s stabilization of a slow phase in HMM but not S1 (Fig. 3). Mavacamten also stabilizes this SRX-like, auto-inhibited biochemical state at physiological ionic strengths (Fig. 5).

**Fig. S2.**
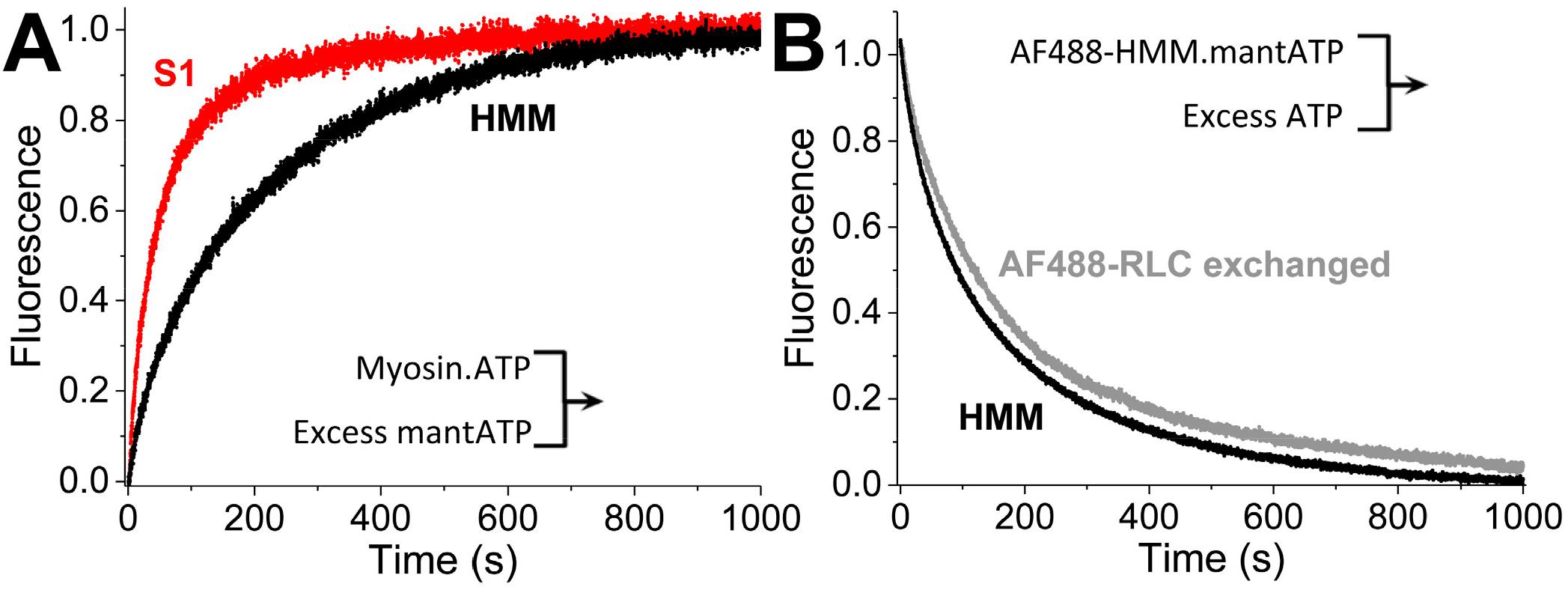
Basal ATP turnover controls: Myosin.ATP mixed with mant-ATP; basal turnover with exchanged light chain. (A) To ensure the mant-nucleotide was not inducing non-physiological phenomenon in the basal mant-turnover experiments, we performed the inverse mix: incubating 0.4 μM S1 or 0.2 μM HMM with 4.0 μM ATP and then mixed with 400 μM mant-ATP. HMM remains biphasic, unlike S1. (B) Fluorescently-labeled and exchanged-on RLC, used in (TR)^2^FRET experiments, did not disrupt the SRX-like basal ATP turnover of HMM. The same bi-phasic behavior was observed. Fits reported in Table S1.

**Fig. S3.**
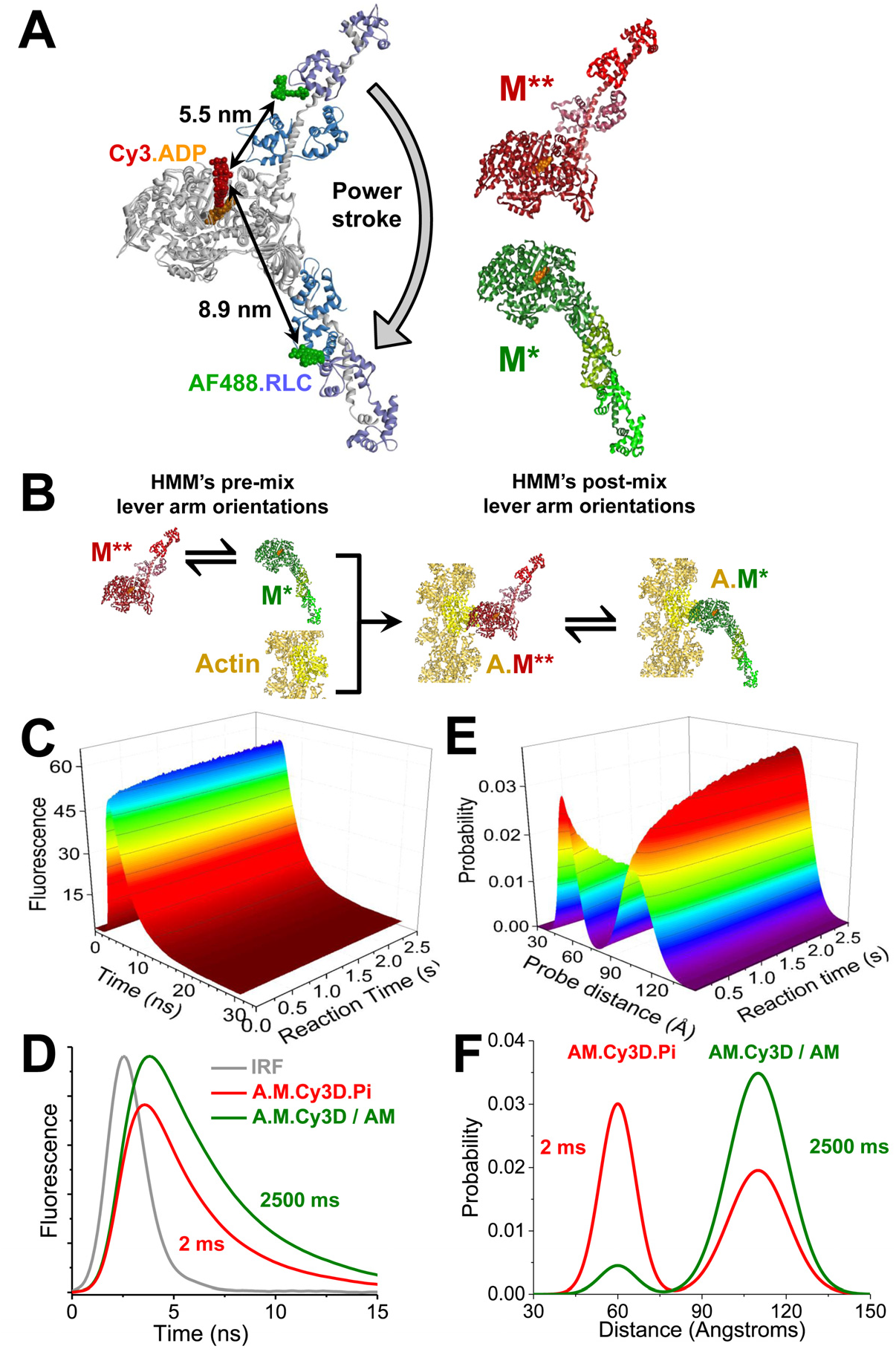
Light-chain domain rotation data acquisition and analysis with (TR)^2^FRET. (A) Fluorescently-labeled HMM, dyes modeled on perpendicular to the protein surface, PDB: 1B7T, 1DFL. Donor fluorophore is AlexaFluor488-RLC and acceptor fluorophore is Cy3-ATP. Distances represent dye center-dye center. Powerstroke occurs from the lever-up pre-powerstroke state (red myosin S1 cartoon, lever up) to the lever-down post-powerstroke state (green myosin S1 cartoon, lever down). S1 depicted for simplicity, HMM used. (B) Stopped-flow mix of myosin with actin, to detect FRET between the lever arm and catalytic domain of cardiac HMM. Equilibrium of structural states of myosin with fluorescently-labeled and exchanged-on RLC with excess fluorescent Cy3-ATP, mixed with actin, detecting the depletion of the lever-up M** structural state (red). (C) 2000 time-resolved FRET waveforms were acquired with 125 picosecond resolution. Each waveform results from after excitation with a 5000 Hz pulsed laser, acquired following stopped-flow mixing of 0.1 μM AF488-labeled cardiac HMM equilibrated with 2.0 μM Cy3-ATP, mixed with 20 mM Actin containing 1.0 mM ATP, depicting a total integrated fluorescence intensity change of 27% for the DMSO control shown. (D) The first and last time-resolved fluorescent waveforms shown in C. Laser pulse (IRF) is shown in gray. (E) Global two-distance fit to each acquired waveforms in C, depicting a mole fraction change of 0.49 for DMSO. Every waveform detected in C was fit to this two-distance model (SI Methods). (F) First and last two-distance fit shown in E.

**Fig. S4.**
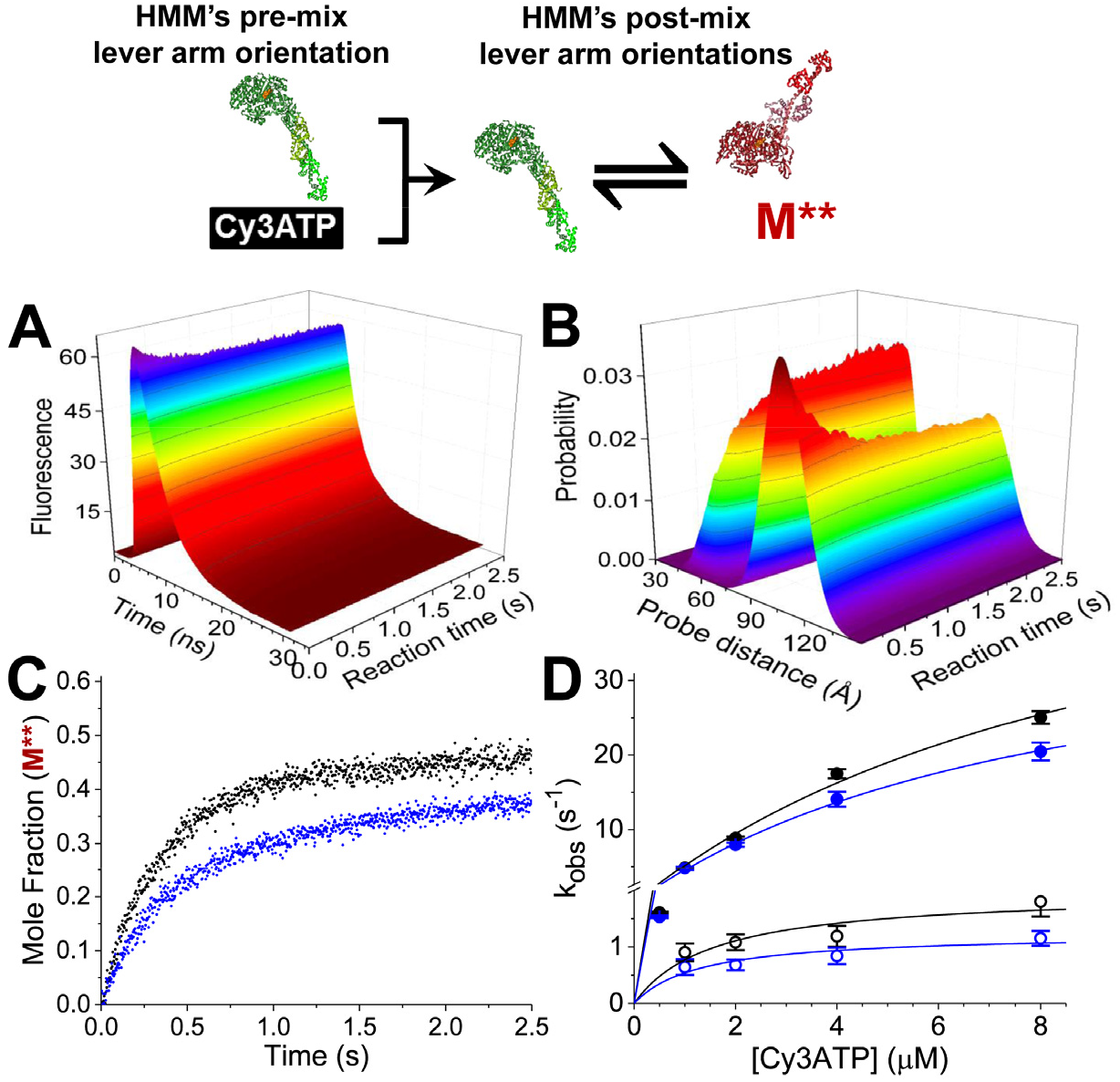
Recovery stroke detected with (TR)^2^FRET. (A) Transient time-resolved FRET waveforms acquired after mixing 0.1 μM AF488 labeled cardiac HMM with 2.0 μM Cy3-ATP, depicting a total integrated fluorescence intensity change of 25% for DMSO. Reaction mixed represented above; S1 shown for simplicity. (B) Global two-distance structural state model was fit to each acquired waveforms in A, depicting a mole fraction change of 0.46. (C) Plotted mole fraction of the M**, lever-up, post-recovery stroke structural state at 2.0 μM Cy3ATP in the absence (black) or presence (blue) of 10 μM Myk461. (D) Observed rate constant for the recovery stroke over a range of nucleotide concentrations. k_obs1,DMSO_ = 58.5s^−1^[ATP]/(10.4μM+[ATP]), k_obs1,Myk461_ = 42.1s^−1^[ATP]/ (8.3μM+[ATP]), k_obs2,DMSO_ = 1.9s^−1^[ATP]/(1.6μM+[ATP]), k_obs2,Myk461_ = 1.2s^−1^[ATP]/(1.3μM+[ATP]).

**Fig. S5.**
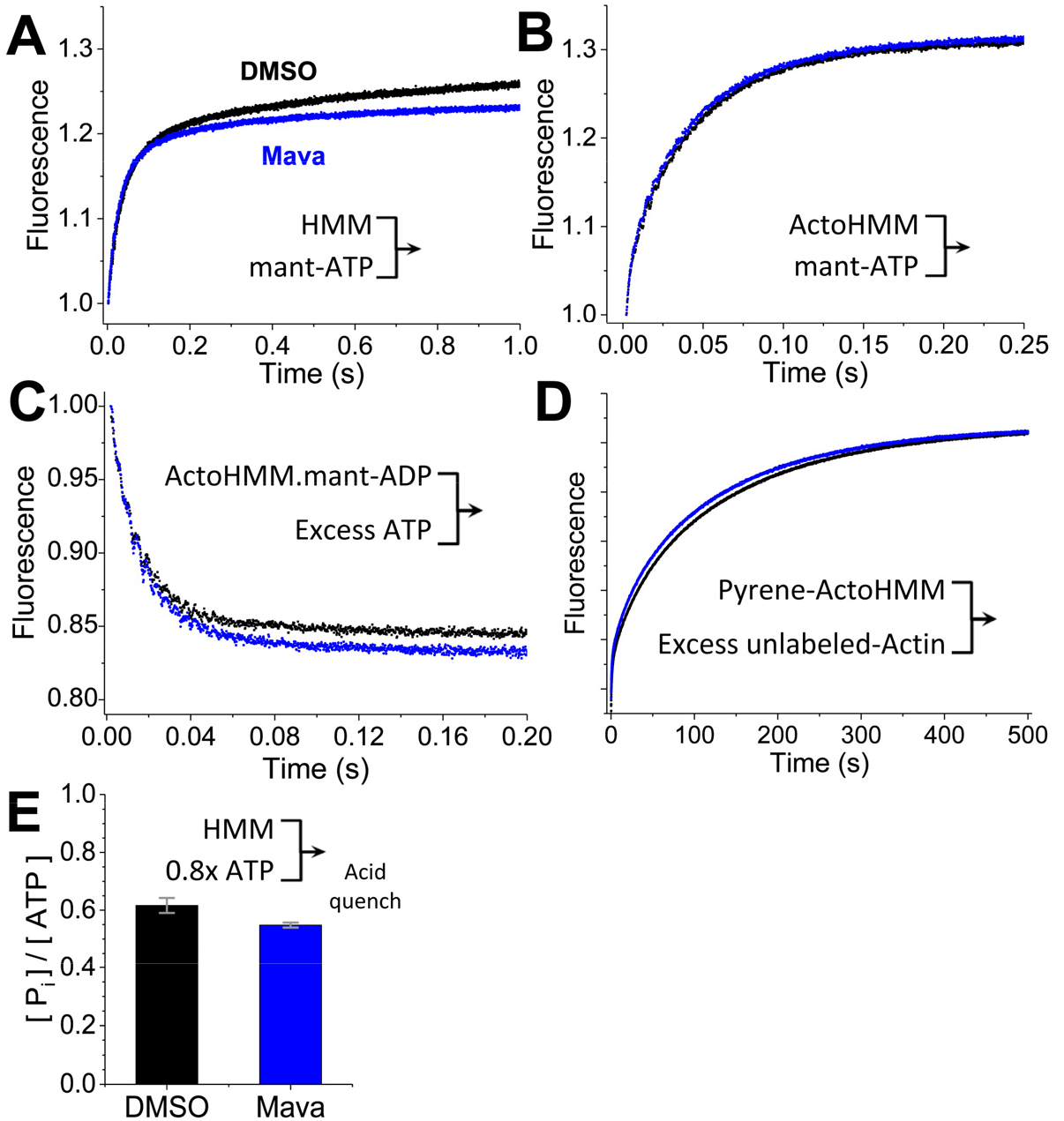
Mavacamten does not change the kinetics of several steps in the myosin ATPase cycle. (A) Mant-ATP binding to HMM. 2.0 μM mant-ATP binding to 0.1 μM HMM in the presence of 1% DMSO (black) or 5.0 μM Myk461 dissolved in DMSO (blue). (B) Mant-ATP binding actoHMM. 2.0 μM mant-ATP binding pre-incubated 5.0 μM actin and 0.1 μM HMM. Observed rate constants given in Table S1. Individual, representative mixes with transients from the average of 6-10 shots of the stopped flow. Two exponential fit and error bars shown in Table S1 from n = 6 mixes, ± SEM. (C) ADP dissociation detected with 5.0 μM actin, 0.1 μM HMM and 2.0 μM mant-ADP chased with 2.0 mM non-fluorescent ATP. Black is DMSO control, blue is 5.0 μM Myk461. Observed rate constants reported in Table S1, ± SEM from n = 6 mixes. (D) Actomyosin dissociation detected with incubation of 1.5 μM pyrene-actin and 0.4 μM HMM mixed with 20 μM unlabeled-actin. Black trace is DMSO, blue is 5.0 μM Myk461. Observed rate constants reported in Table S1 ± SEM from n =6. (E) Hydrolysis detected with malachite green following acid quench with perchloric acid. 12.5 μM two-headed HMM was mixed with 20 μM ATP (5:4 stoichiometry) and incubated for 5.0 s, then quenched with 0.6 M perchloric acid and detected with malachite green. The concentration of P_i_ was determined with a standard curve.

